# Analysis of tumor-derived and cross-presented peptide antigens defines improved immunotherapeutic strategies

**DOI:** 10.64898/2026.02.23.707477

**Authors:** Yufei Cui, Kien Phuong, Heidi Temple, Amy J. Wisdom, Nouran S. Abdelfattah, Stefani Spranger, Forest White

## Abstract

**Background:** Cross-presentation of tumor antigens by antigen-presenting cells (APCs) is essential for initiating effective anti-tumor T cell immunity. The presence of cross-presenting immune cells across multiple solid tumors correlates with improved clinical outcomes. Despite the importance of this process, the identities and characteristics of tumor-derived MHC-I antigens that are cross-presented by APCs remain largely undefined, limiting rational design of targeted immunotherapies.

**Methods:** We performed an immunopeptidomic analysis of cross-presented glioblastoma (GBM) antigens on APCs, including bone marrow–derived macrophages, bone marrow–derived dendritic cells, and splenic dendritic cells, using SILAC labeling and in vitro co-culture systems. Additionally, we also profiled endogenous APC and tumor antigen repertoires. We made selected cross-presented antigen targets into mRNA vaccines and evaluated their immunogenicity in comparison to tumor endogenous antigens *in vivo*.

**Results:** We identified over one thousand putative cross-presented GBM antigens. Comparative analysis of endogenous APC and tumor antigen repertoires revealed that cross-presented antigens possess distinct features and are predominantly shaped by intrinsic antigen processing and presentation pathways within APCs, resulting in limited cross-presentation of tumor-specific epitopes. Two doses of mRNA encoding cross-presented tumor-specific epitopes delayed tumor growth and elicited robust antigen-specific T cell responses.

**Conclusion:** Our findings define the landscape and constraints of tumor antigen cross-presentation in GBM and establish a framework for improved antigen selection in the development of next-generation GBM immunotherapies.

## Background

GBM is one of the deadliest human tumors, with less than a 5% five-year survival rate. GBM tumors are characterized by diffuse infiltration in parenchymal normal brain tissue, high inter-and intra-tumoral heterogeneity, and resistance to conventional therapies including radiotherapy and temozolomide chemotherapy^1,2^. Although immunotherapies have demonstrated favorable response in some solid tumors, GBM remains unresponsive to checkpoint blockade and other targeted immunotherapies^3–5^. This limited immunotherapeutic efficacy is largely due to a profoundly immunosuppressive tumor microenvironment, marked by dysfunctional T-cell activity, extensive myeloid infiltration, and impaired antigen presentation^6^. As a result, there is increasing interest in alternative immunotherapeutic strategies to enhance T-cell priming through optimized antigen delivery to synergize with immune checkpoint inhibitor treatments and improve treatment efficacy. Central to these efforts is a better understanding of the antigenic landscape that can be recognized by cytotoxic T lymphocytes and result in immune activation against tumor cells.

Antigen cross-presentation (XPT) is a key biological process that enables the initiation of anti-tumor CTL responses^7,8^. In this process, professional antigen-presenting cells (APCs) such as dendritic cells (DCs) and macrophages take up exogenous tumor derived proteins and present processed peptides on MHC class I molecules, priming naïve T cells to recognize tumor antigens^9^. XPT is essential for generating an anti-tumor immune response, even when tumor cells downregulate direct MHC-I presentation. In this context, cross-presenting APCs activate functional cytotoxic T cells that recognize remaining MHC-I positive tumor cells, secrete cytokines to induce or upregulate MHC-I expression on other tumor cells, and activate an NK cell response to MHC-I deficient tumor cells ^10,11^. In GBM, APC-mediated XPT has been shown to occur both in the tumor microenvironment and in draining cervical lymph nodes, where it contributes to activation of tumor-specific CD8 T cells and synergizes with checkpoint blockade^12,13^. However, despite the immunological importance of XPT, the identity of native tumor peptides that are cross-presented by DCs or macrophages in GBM has remained largely undefined.

Most prior work on XPT has relied on model antigens such as OVA or viral epitopes, leaving a major gap in our understanding of tumor antigen XPT. Mass-spectrometry–based immunopeptidomics enables unbiased discovery of naturally presented MHC-I ligands from tumors and APCs^14^. Within GBM, immunopeptidomic profiling has been used to uncover tumor-associated antigens and shared peptide motifs, but the majority of studies focused on endogenous presentation by tumor cells, rather than cross-presentation by professional APCs, which ultimately determines the priming of anti-tumor T-cell responses. To date, only a handful of immunopeptidomic studies have attempted to map cross-presented tumor antigens, and these datasets were limited in scale and resolution^15,16^.

APC-based cancer immunotherapies such as dendritic cell vaccines and engineered macrophage platforms depend on cross-presentation of immunogenic tumor associated antigens to elicit therapeutic efficacy^17,18^. Clinical DC vaccine trials in GBM, ranging from peptide-loaded DCs to tumor lysate–pulsed formulations, have shown encouraging but variable responses, partly due to uncertainty surrounding which tumor antigens were effectively processed and displayed by APCs in vivo that led to effective antitumor responses^19,20^. A comprehensive mapping of XPT antigen repertoire in DCs and macrophages would therefore provide a rational base for designing targeted vaccines that optimize antigen choice, improve T-cell priming, and enable more durable responses in GBM patients.

Here, we use mass-spectrometry–based immunopeptidomics to define, for the first time, the landscape of cross-presented GBM antigens on dendritic cells and macrophages using syngeneic murine CT2A and GL261 glioma models. By comparing to APC and tumor endogenous antigens, we identified different classes of XPT antigens, including those also presented endogenously on tumor cells and/or immune cells, as well as XPT antigens that were uniquely presented only in the context of cross-presentation. The biogenesis pathways for each class were inferred, providing insight into the principles that govern XPT. Finally, we compared the immune response generated by different classes of XPT antigens, demonstrating the relevance of selected classes of XPT antigens to translational research. These findings offer insight into antigen processing bottlenecks in XPT, and highlight actionable peptide targets for improved immunotherapy design for GBM treatment.

## Results

### Identification of cross-presented GBM antigens on primary dendritic cells and macrophages

We analyzed a published dataset of single-cell RNA-sequencing from 44 human glioma tissues to infer potential cross-presentation in the tumor microenvironment^25^. Using myeloid signatures from Abdelfattah et al, we identified the MCsig_8 cell cluster that expresses canonical DC markers and MCsig_1, 2, 6 and 7 clusters that express microglia/macrophage markers (Figure 1a). High levels of XCR1, CLEC9A, BATF3, and IRF8 suggest cDC1 lineage, which specializes in cross-presentation. CXCL9, CXCL10, TAP1, and TAP2 expression were highly expressed in MCsig_6 cluster, indicating an inflammatory macrophage program and potentially activated cross-presentation. Inference on cell-cell communication revealed crosstalk among tumor cells, T cells, DC, and microglia/macrophage clusters (Figure 1a). The inferred ligand–receptor interactions suggest antigen uptake, as well as tumor APC engagement (Supplementary Figure 1a). For instance, CD74–CD44 and CD74–CXCR4 interactions suggest tumor cells interacting with DC/macrophage clusters, promoting APC survival and positioning within antigen-rich tumor niches. C3–CR4 (C3–ITGAX/ITGB2) interactions suggest capture of complement-opsonized tumor antigens by DC-like cells. Furthermore, GAS6–AXL and GAS6–MERTK signaling between myeloid subsets indicate efferocytic uptake of apoptotic tumor cells, supporting cross-presentation of tumor antigens. Together, this evidence suggests that APCs within GBM tumors, including DCs and macrophages, are molecularly equipped to cross-present tumor antigens.

**Figure 1.**
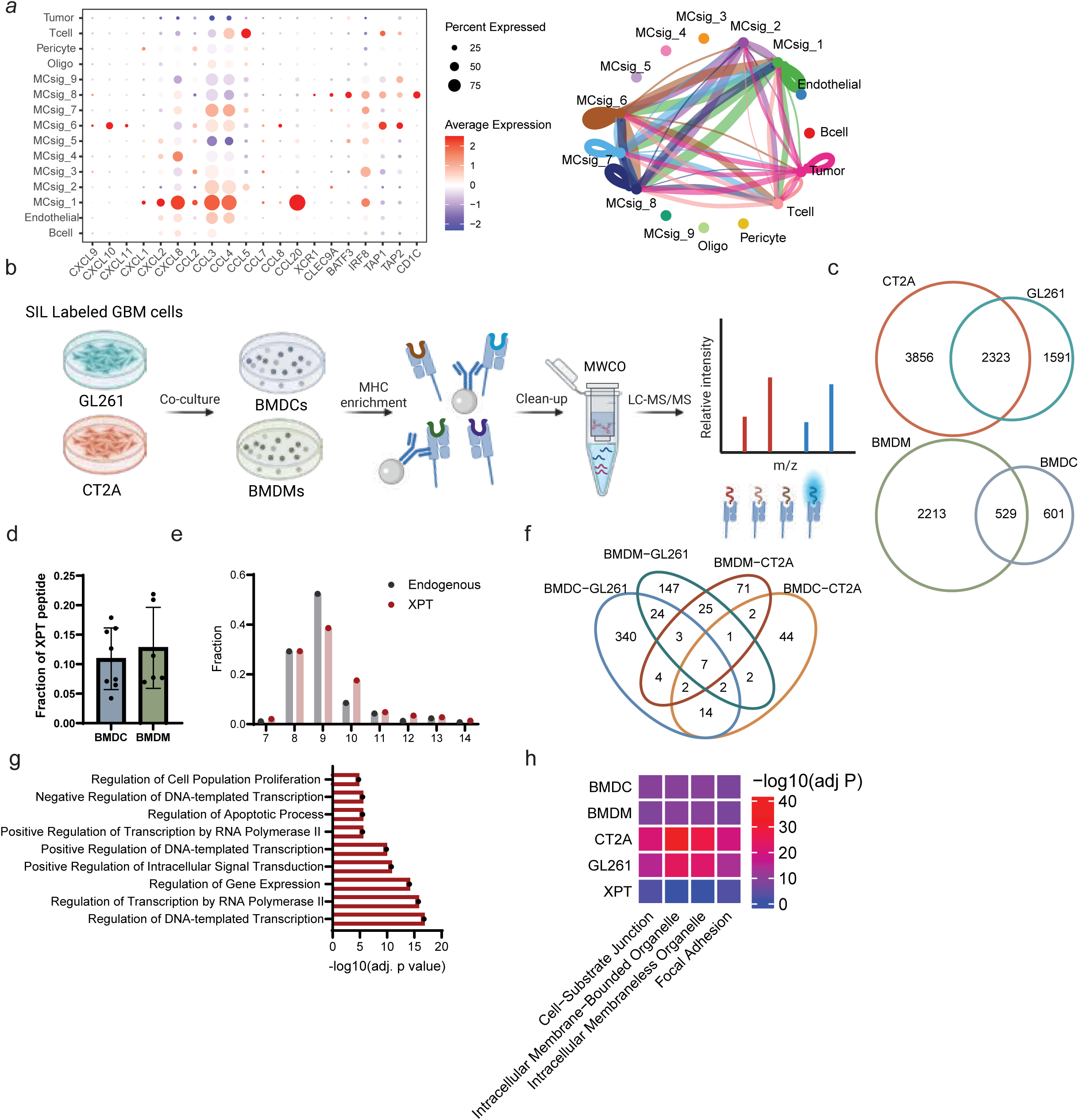
Identification of XPT GBM antigens from BMDMs and BMDCs. a). ScRNA seq analysis of cross-presenting immune cells in human GBM. (Left) Average gene expression of chemokine and cDC1 lineage genes in different cell types from ScRNA seq analysis. Msig_8 representing DC cluster show high expression of cDC1 lineage genes. (right) Cellchat inferred interactions among DC, macrophage clusters, tumor cells, and T cells. b). Schematic of immunopeptidomics analysis of SILAC labeled tumor cells and primary immune cells co-culture to identify XPT peptides. c). Venn diagram of overlap between MHC-I peptide repertoire of GBM cell lines GL261 and CT2A, and between APCs BMDM and BMDC. d). Fraction of XPT peptides in the total peptides profiled from cross-presenting BMDM and BMDC. p value calculated using student’s t-test, where the variance difference between two groups is determined by F-test. No significant difference was observed between two groups (p>0.05). e). Sequence length distribution of cross-presented and endogenous peptide. Amino acid sequence length between 7 and 14 were shown. f). Comparison of XPT antigens identified from BMDM or BMDCs cross-presenting GL261 or CT2A derived antigens. g). GO biological process enrichment analysis of source proteins from XPT antigens on BMDCs and BMDMs. Total profiled tumor endogenous peptides associated source proteins were used as enrichment background. P-value was adjusted by Benjamini-Hochberg method. Significant pathways (p<0.05) were colored in red. h). Heatmap of GO cellular compartment enrichment analysis of source proteins from XPT antigens, BMDC endogenous antigens, BMDM endogenous antigens, CT2A endogenous antigens, and GL261 endogenous antigens. Whole proteome was used as enrichment background. P-value was adjusted by Benjamini-Hochberg method. Color bar showed -log10 adjusted p-value.

To unbiasedly profile XPT MHC-I antigens on APCs, we established a co-culture system of GBM tumor cells and primary APCs, including bone marrow derived macrophage (BMDMs) and bone marrow derived dendritic cells (BMDCs) (Figure 1b). Murine GBM cell lines GL261 and CT2A were engineered using CRISPR knockout of MHC-I expression as previously described to eliminate confounding tumor endogenous MHC-I peptides^21^. MHC-I null tumor cells were cultured in stable isotope labeled (SIL) amino acid containing media for 72hrs to allow incorporation of the SIL amino acids into tumor derived proteins (Supplement Figure 1b). SIL amino acids tyrosine, phenylalanine, and asparagine were selected because they are among the anchoring residues of tumor cell MHC-I peptides, and are present in 80% of MHC-I peptides (Supplementary Figure 1c). Global protein expression profiling was used to quantify the labeling efficiency of tumor proteins after 72hrs; this analysis revealed that over 94% of peptides fully incorporated the SIL amino acids (Supplementary Figure 1d). Differentiated BMDMs or BMDCs were subsequently co-cultured with labeled GBM tumor cells for 18hrs followed by mass spectrometry based immunopeptidomic analysis (e.g., MHC-I enrichment, peptide elution, and liquid chromatography tandem mass spectrometry (LC-MS/MS) analysis). Acquired MS/MS spectra were searched with SIL amino acids as dynamic modifications, and peptides containing SIL amino acids were identified as putative XPT peptides. MHC-I peptides from parental, MHC-I expressing GBM tumor cells and APC monocultures were also collected from multiple replicates to compare with XPT peptides. Overall, we identified 7770 peptides from GBM tumor cells, 3343 peptides from APCs, and 688 XPT peptides. Both tumor cell types (CT2A and GL261) and APCs (BMDMs and BMDCs) naturally presented different MHC-I peptide repertoires (Figure 1c). In BMDMs and BMDCs, approximately 10-12% of MHC-I peptides were putative XPT antigens (Figure 1d). Profiled XPT peptides shared similar length distribution with endogenous APC MHC-I peptides, but XPT peptides showed a slight preference for longer peptides^26^ (Figure 1e). Gibbs clustering analysis revealed that unlike tumor or APC endogenous MHC-I peptides, XPT peptides did not clearly represent H2-Kb or H2-Db binding motifs (Supplementary Figure 1e). Correspondingly, XPT antigens displayed lower binding affinity to the MHC-I alleles compared to tumor or APC endogenous peptides, suggesting decreased peptide binding affinity restriction during cross-presentation (Supplementary Figure 1f).

Comparing cross-presented peptides from BMDMs and BMDCs, we found that the identities of XPT peptides were dependent on the type of APC and the tumor cell type. There was limited overlap among XPT peptides between BMDMs and BMDCs cross-presenting GL261 or CT2A antigens, and only 7 peptides were detected in all four scenarios (Figure 1f, Supplementary Figure 1g). To understand the internal biological processes represented by MHC-I peptide presentation, we performed biological process enrichment analysis on source proteins from antigens presented by APCs, GBM tumor cells, or XPT. Some of the top enriched pathways for GL261, CT2A, and BMDC cell endogenous antigens were shared, including pathways associated with cytoplasmic translation, protein transport, and macromolecule biosynthetic organization, whereas BMDMs specifically enriched for protein modification associated processes (Supplementary Figure 1h). However, XPT antigens enriched for biological processes different from any endogenous protein repertoires, representing axonogenesis, gene regulation and neurogenesis. Using all presented proteins from APCs as a background, XPT antigens enriched for biological processes associated with transcription regulation, a process that is often upregulated in tumors^27^ (Figure 1g). We wondered if the difference in enriched biological process could be due to differential sub-cellular localization. However, XPT antigens enriched for similar cell compartments compared to APC or tumor endogenous antigens, both representing cell substrate junction and intracellular organelles (Figure 1h).

Since BMDMs and BMDCs displayed different XPT antigens, we repeated the cross-presentation experiment using splenic DCs in co-culture with MHC-I knock-out GL261 tumor cells to include another more physiologically relevant type of APC for priming an anti-tumor response. Unlike DMXAA-matured BMDCs enriched for cDC1s, splenic DCs contain a more balanced ratio of cDC1s and cDC2s^28^ compared to BMDCs. Approximately half of the endogenous peptides from splenic DCs were shared with BMDC or BMDMs (Supplementary Figure 2a). SIL amino acids were similarly represented on splenic DC MHC-I peptide sequences compared to BMDCs and BMDMs (Supplementary Figure 2b). We profiled a slightly increased level (approximately 18%) of XPT antigens in splenic DCs compared to BMDMs or BMDCs (Supplementary Figure 2c). In total, 445 XPT peptides from BMDCs, 290 from BMDMs, and 567 from splenic DCs were detected (Supplementary Figure 2d). Splenic DCs cross-presented a different repertoire of peptides, in which only 11.1% peptides and 15.7% source proteins are shared with BMDMs or BMDCs (Supplementary Figure 2e). Even though the source proteins are different between the XPT antigens on the different APC types, XPT antigens from splenic DCs still enriched for cellular compartments consistent with XPT antigens from BMDMs and BMDCs, including cytoskeleton and cell substrate junctions (Supplementary Figure 2f). Collectively, we profiled 1192 XPT peptides from three types of APCs, representing 1066 proteins.

We compared the sequences of total XPT peptides with endogenous peptides from source GBM cells or APCs to define 3 groups of XPT peptides (Figure 2a, Supplementary Table 7-9). Categorization of XPT peptides was based on their relationship with GBM cell or APC endogenous presentation: ‘XPT-shared’ peptides that were endogenously presented on APCs and tumor cells, ‘XPT-only’ peptides that were not detected in either APC or GBM endogenous repertoire, and ‘XPT-tumor’ peptides that were also presented endogenously on tumor cells but not presented endogenously on APCs. We reasoned that in the cross-priming process, XPT-tumor peptides could play an important role because they can enable T cell recognition of the tumor. We hypothesized that the XPT-shared antigens might augment the immune response but might be subjected to immune tolerance. On the other hand, XPT-only peptides might not lead to an anti-tumor immune response as they are only cross-presented on APCs and not presented on the tumor cells. Intriguingly, in the XPT repertoire, XPT-tumor peptides were the most infrequent, with only ∼5% (53 of the 1192) of XPT peptides classified as XPT-tumor, while ∼15% (179 of the 1192) of XPT peptides were XPT-shared, and ∼80% (960 of the 1192) of XPT peptides were XPT-only (Figure 2b).

**Figure 2.**
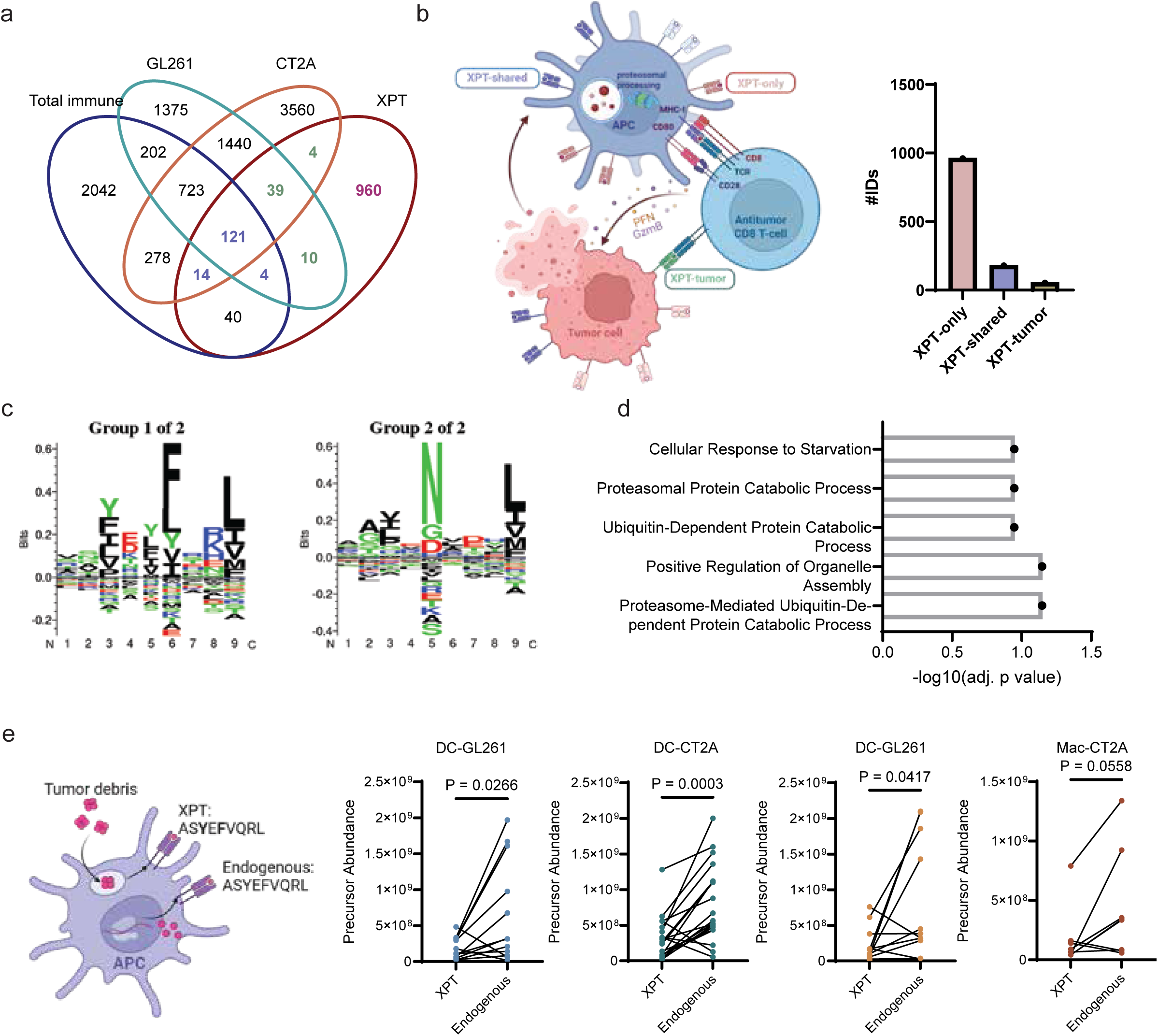
XPT-shared antigens are enriched for intracellular membrane bounded organelles. a). Venn diagram of overlaps among endogenous antigens on total immune cells presented, endogenous antigens on GL261 or CT2A tumor cells, and XPT antigens. b). Number of peptides in three categories of XPT antigens: XPT-only, XPT-shared, and XPT-tumor. c). Sequence motif enrichment of XPT-shared antigens corresponding to H2-Kb motif (left) and H2-Db motif (right). d). GO biological process enrichment analysis of source proteins from XPT-shared antigens. Total XPT antigens associated source protein repertoire was used as enrichment background. P-value was adjusted by Benjamini-Hochberg method. Significant pathways (p<0.05) were colored in red. e). Precursor abundance (Integrated MS1 intensity) comparison between XPT-shared peptides and corresponding endogenous peptides in BMDMs or BMDCs cross-presenting GL261 or CT2A antigens.

### XPT-shared peptides positively correlate with endogenous presentation on APCs but not tumor cells

To gain a better understanding of the XPT peptides, we first focused on the XPT-shared antigens: XPT peptides that were also endogenously presented on tumor cells and APCs. XPT-shared antigens were enriched for H2-Kb and H2-Db binding motifs (Figure 2c). Using the total XPT source proteins as background, we found that XPT-shared peptides were enriched, although not significantly, for ubiquitin dependent protein catabolic processes that are features of the protein degradation pathway for MHC-I peptide presentation (Figure 2d). Two potential routes for processing of XPT peptides have been proposed: a canonical, cytosolic route typically used for processing of endogenous proteins and peptides, and a vacuolar route that is largely TAP-independent^29,30^. Our data suggests that the group of XPT-shared antigens were most likely processed via canonical antigen processing pathways, as they have similar specificity to H2 alleles compared to APC endogenous peptides, and enrich for proteins associated with proteasomal processing pathways. To assess the relative presentation level of XPT-shared peptides to their endogenous counterparts, we compared the MS precursor abundances of a subset of XPT peptides with their endogenous analogs. Endogenous peptides were presented at significantly higher level compared to XPT antigens (Figure 2e). To ensure that this result was not due to run-to-run variability we compared the abundance ranks of peptides across different runs and showed that the abundance rank of each peptide was comparable (Supplementary Figure 3a). Abundance rankings between the XPT-shared antigens matched to the rankings of the corresponding endogenous peptides, suggesting that processing of these XPT-shared peptides may be similar to endogenous antigen processing in APCs. (Supplementary Figure 3b).

To more accurately quantify the correlation between endogenous presentation and cross-presentation of the XPT-shared sequences, we used SureQuant targeted mass spectrometry to quantify the presentation amounts of 11 selected XPT-shared peptides ^23^ (Figure 3a, Supplementary Table 3). Heavy synthetic peptides with different SIL amino acids than the XPT SIL amino acids were folded into recombinant H2-Kb or H2-Db monomers and then spiked into GL261-BMDM or BMDC co-cultures or each monoculture during MHC-I enrichment. In SureQuant, matching of the SIL peptide MS2 to pseudospectra triggered high-resolution targeted MS2 scans at the corresponding m/z of the endogenous or XPT versions of the same peptide (Supplementary Table 2). The workflow allowed for endogenous and XPT peptide sequence validation via retention time and peptide spectral matching to the synthetic standard. To estimate the presented copies per cell of each peptide, the ratio of the targeted product ion abundances between the synthetic peptide and endogenous or XPT peptides was used. All 11 XPT-shared peptides were validated to be presented endogenously on APCs and GL261 cells, and were cross-presented at detectable levels (Supplementary Figure 4). The selected peptides were endogenously presented by GL261 cells or APCs at hundreds to thousands of copies/cell, and cross-presented at tens to low hundreds of copies per cell (Supplementary Figure 5). This lower level of cross-presentation is consistent with previous reports that direct presentation of viral antigens from infected cells outnumber the corresponding cross-presented antigens^31,32^. Some of the selected peptides, such as Dync1h1 and Elp2 peptides, were presented at high levels on GL261 tumor cells but endogenously presented on APCs and cross-presented at lower levels. Alternatively, some peptides were presented on tumor cells and APCs at similar amounts. Interestingly, a subset of antigens including H2-D1 and Sgk1 were barely presented on tumor cells, but cross-presented at high levels on the immune cells. Consistent with results from label free quantification, cross-presentation of XPT-shared antigens occurred at a lower level compared to APC endogenous presentation. To assess whether XPT-shared peptides were processed and presented similarly to endogenous tumor-presented peptide or to endogenous APC-presented peptides, we quantified XPT-shared peptide abundance to copies/cell and correlated tumor or APC endogenous presentation with XPT. Strong positive correlation was observed between APC endogenous presentation with cross-presentation, but cross-presentation was not correlated at all with tumor endogenous presentation (Figure 3b). This phenomenon suggests that cross-presentation of XPT-shared antigens is dictated by immune cell endogenous processing and presentation pathways, likely through the cytosolic XPT pathways. BMDCs on average cross-present higher levels of XPT-shared antigens compared to BMDM, although the differences were not significant (Figure 3c). Our quantification indicates that antigens presented at high levels by the immune cells are more efficiently cross-presented, and that antigen presentation preference is distinctly different between APCs and tumor cells. This result highlights the importance of understanding antigen processing and presentation rules in APCs in designing immune stimulatory therapies, in addition to current antigen discovery efforts that primarily focus on profiling tumor presented antigens.

**Figure 3.**
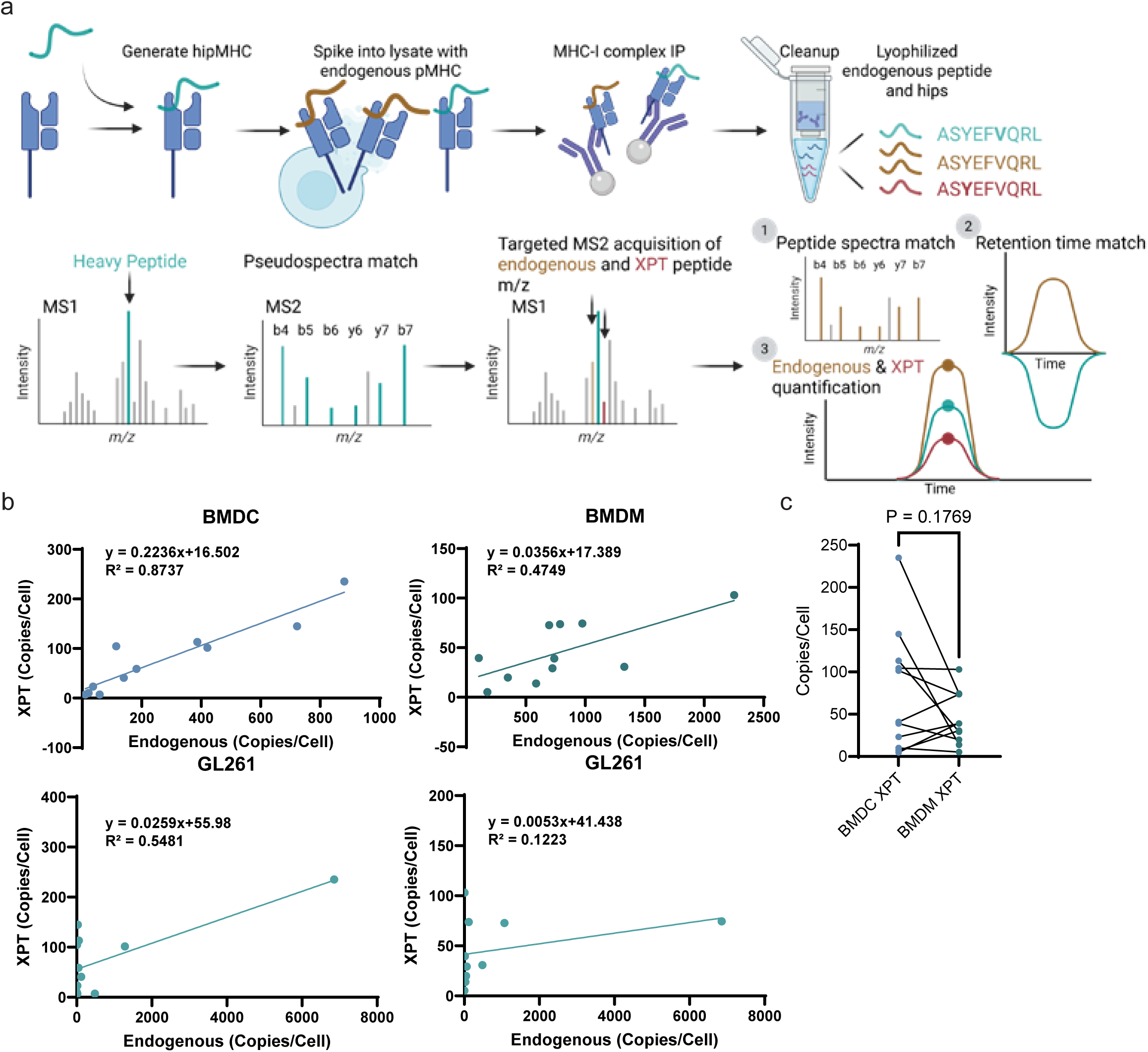
Quantitative analysis reveals positive correlation between XPT and endogenous antigen presentation on APCs. a). Schematic of SureQuant workflow. Stable isotope labeled heavy peptide are folded into empty MHC-I complexes and spiked into lysate. MS2 scan of heavy peptide was used to trigger MS2 scan of endogenous light peptide or XPT peptide with SILAC labeling at corresponding m/z offset. b). Correlation of copies/cell endogenous peptide presentation of APCs or GBM tumor cells with cross-presentation level. Simple linear regression was performed to interpolate slope and intercept. c). Copies per cell cross-presentation level comparison between BMDM and BMDC of selected XPT antigens. Paired t-test was performed to compare between two groups.

### XPT-tumor peptides are less favored during cross-presentation and are enriched in mitochondrial proteins

XPT-tumor peptides, although expected to be most important class of XPT peptides for priming of an anti-tumor response, comprise the smallest portion of total XPT peptides (4.7%). Peptides presented only on tumor cells are less likely to be cross-presented compared to peptides commonly presented on both tumor and APCs (Figure 4a). Indeed, only 49 peptides from GL261 and 43 from CT2A were cross-presented, comprising only 1.7% of endogenous peptides specific to tumors. On the other hand, of the endogenous peptides presented on both tumors and APCs, 11.9% were also cross presented. This difference again suggests that antigen processing and presentation is different in APCs from tumor cells; as a result, tumor endogenous peptides are less likely to be cross-presented on APCs. Intriguingly, even though XPT-tumor peptides appeared to be disfavored among the overall cohort of XPT peptides, most (81%) were shared between GL261 and CT2A, with only 4 peptides specific to CT2A and 10 to GL261 (Supplementary Figure 6a), suggesting some selective mechanism as to which tumor peptides were cross-presented on APCs. XPT-tumor peptides mapped to 26 source proteins, with 16 of these shared between GL261 and CT2A, while 4 were specific to GL261 and 6 were specific to CT2A. We selected 20 peptides from the XPT-tumor peptides to validate the cell type specificities with targeted PRM analysis of GL261, CT2A, BMDM and BMDC monocultures, and confirmed that 19/20 are presented only on tumor cells and not detectable on either BMDMs or BMDCs (Supplementary Figure 6b). Given that the initial analyses were all performed by data-dependent analysis (DDA) that is known to have analysis-to-analysis variability and is not comprehensive, this PRM validation result suggests that our general classification of XPT peptides into different categories based on tumor and APC endogenous peptide repertoire was accurate. Compared to the total XPT repertoire, XPT-tumor peptides displayed lower binding affinity, but higher than randomly sampled peptides from the proteome with the same length distribution (Figure 4b). There was no significant difference between binding affinities of CT2A and GL261 XPT-tumor peptides. It is also worth noting that the percentage of NetMHC Pan predicted MHC-I binders (EL_Rank<2) are similar across CT2A XPT peptides, GL261 XPT peptides, and total XPT peptides for either H2-Kb or H2-Db allele. We found that a high ratio of CT2A and GL261 XPT-tumor peptides (39.5% and 40.8%) were derived from a single source protein Vimentin (Figure 4c). Vimentin is a cytoskeletal protein overexpressed in many types of tumors, including GBM, and has been shown to be a therapeutic target^33–35^. Interestingly, none of the peptides were predicted binders of H2-Kb or H2-Db alleles (Supplementary Figure 6c). Other XPT-tumor peptides mostly mapped to different source proteins, with only Atp5pf, Psmc1, Psmc4, and Fau having multiple XPT-tumor peptides. We wondered if these source proteins were over-represented in tumor endogenous peptides, which could explain their favored cross-presentation on APCs. Indeed, in GL261 and CT2A endogenous peptide repertoires, these source proteins had the most mapped peptides (Supplementary Figure 6d). However, except Vimentin, these source proteins with multiple XPT-tumor peptides were not the most highly expressed proteins in GBM tumor cells, as determined by global protein expression profiling (Supplementary Figure 6e). This result suggests that XPT-tumor peptides were potentially derived from proteins favorably presented by tumor cells, even though presentation of these peptides did not correlate with expression abundance^36^. Additionally, XPT-tumor peptides were not limited to source proteins with multiple endogenously presented epitopes. The fractions of XPT-tumor peptides in the tumor endogenous peptide repertoire were variable across each source protein, ranging from 7% to 100% (Figure 4d).

**Figure 4.**
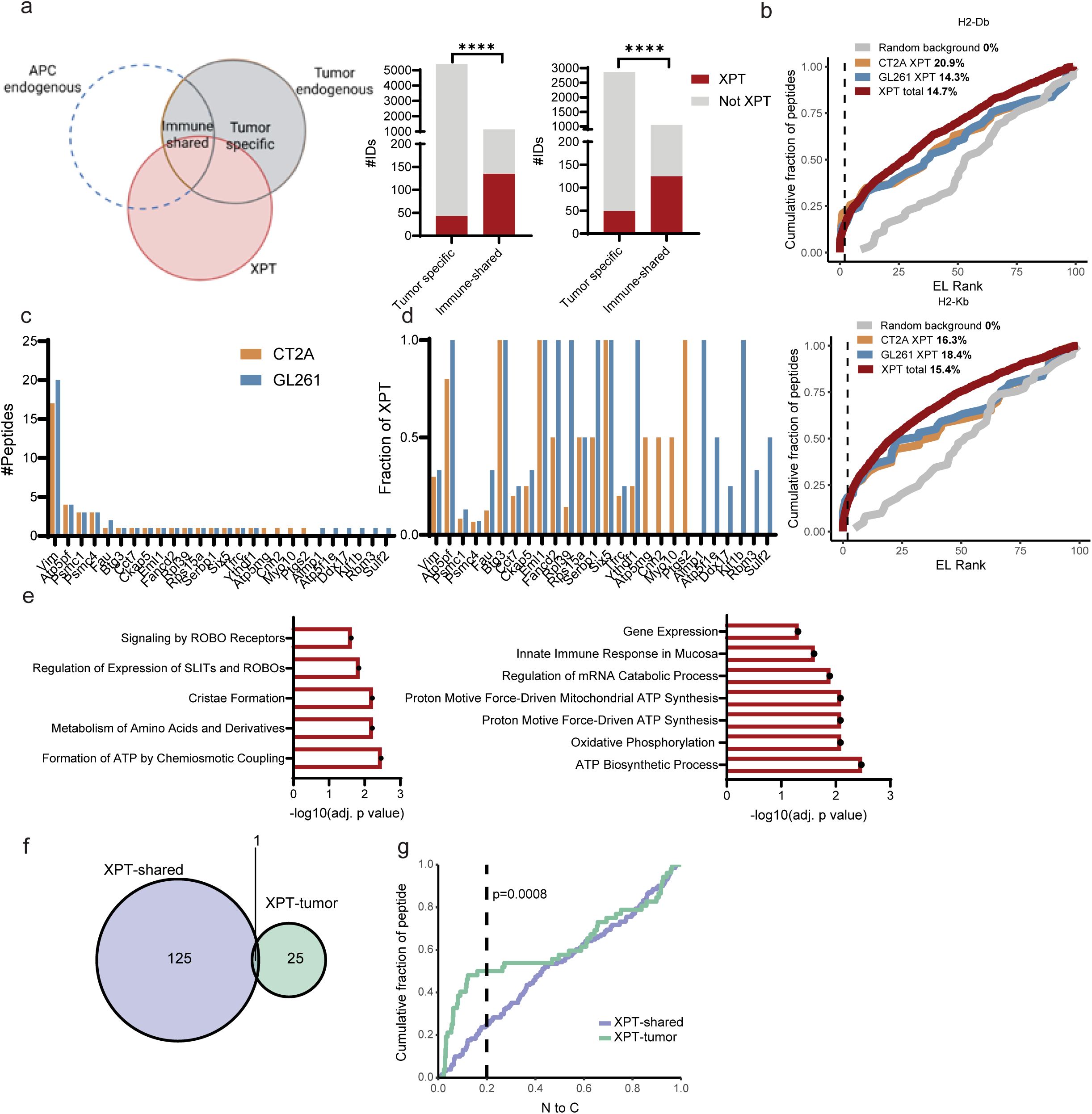
XPT-tumor antigens are less favored during cross-presentation. a). Number of XPT antigens identified in tumor endogenous peptide repertoire or shared peptide repertoire with immune cells. Fisher’s exact test was used to compare difference in ratios of XPT antigens. ****: p<0.0001. b). Cumulative distribution function plot of netMHCPan predicted EL rank of total XPT peptides (red), CT2A XPT-tumor peptides (orange), GL261 XPT-tumor peptides (blue), and random peptides sampled from the proteome (grey) binding to H2-Kb (left) or H2-Db (right). Black line represent EL rank=2. c). Number of XPT-tumor peptides associated with each source proteins d). The fraction of peptides that are cross-presented from XPT-tumor peptide source proteins. e). Reactome pathway (left) and GO biological process (right) enrichment analysis of source proteins from XPT-tumor antigens. Total XPT antigens associated source protein repertoire was used as enrichment background. P-value was adjusted by Benjamini-Hochberg method. Significant pathways (p<0.05) were colored in red. f). Venn diagram of XPT-shared peptide and XPT-tumor source proteins. g). Cumulative distribution function plot of relative position of XPT-tumor peptides (green) and XPT-shared peptides (purple) within the source protein. Position was calculated using the position of the starting amino acid of the peptide. 0 represents N terminus and 1 represents C terminus. Fisher’s exact test was used to compare difference in ratios of XPT-tumor and shared -XPT peptides starting <20% from the N-terminus.

The source proteins of XPT-tumor antigens were enriched for ATP formation pathways and biological processes in the background of the total XPT repertoire, suggesting preference of mitochondria associated proteins during cross-presentation to generate XPT-tumor peptide sequences (Figure 4e). Multiple ATP biogenesis associated proteins including Atp5pf, Atp5mg, and Atp5f1e are cross-presented as XPT-tumor peptides, which were not found in other XPT antigens. Surprisingly, except Vimentin, there was no overlap between source proteins of XPT-shared antigens and XPT-tumor antigens, suggesting potentially different antigen processing pathways that give rise to the two types of antigens (Figure 4f). This hypothesis was also supported by different cellular comparts of source proteins, where the XPT-tumor antigen enriched for cytoskeletal proteins, while the XPT-shared antigens enriched for intracellular membrane-bounded organelle and nucleus (Supplementary Figure 6f). We then asked if the XPT-tumor peptides were derived from different regions within the source protein compared to the XPT-shared antigens, which could indicate differential antigen cleavage during processing. We found that XPT-tumor peptides localized more at the N terminal (<20% from the N terminal) of the proteins, compared to XPT-shared peptides that disfavored the N and C terminus (Figure 4g). We specifically observed a strong preference of XPT-tumor peptides to be at the N terminus of the source protein, potentially suggesting that these XPT-tumor antigens are processed via different XPT pathways compared to XPT-shared antigens (vacuolar vs. cytosolic), giving rise to different source protein preferences and sequence preferences. Biophysical properties of the XPT-tumor peptides are more similar to tumor endogenous peptides than APC endogenous peptides, including higher GRAVY score, isoelectric point, boman index, and lower aliphatic index (Supplementary Figure 6g). These properties explain the suboptimal binding of XPT-tumor antigens to MHC-I molecules, and suggest limited TAP editing and trimming compared to canonical MHC-I peptides.

### XPT-only peptides are enriched for proteins from basement membrane

‘XPT-only’ peptides, those detected only in the cross-presentation setting and not detected in the endogenous tumor or immune cell antigen repertoires, represented the largest subset (∼80%) of XPT peptides. Similar to XPT-tumor antigens, XPT-only antigens have lower binding affinity to H2-Kb and H2-Db allele compared to the total XPT repertoire (Figure 5a). However, the distribution of XPT-only peptides within the source proteins was similar to that of XPT-shared peptides (Figure 5b). We used enrichment analysis to determine the biological processes and cellular compartments associated with these proteins. Source proteins for XPT-only peptides were enriched for signal transduction and nervous system development processes, suggesting their origin from GBM tumor cells (Figure 5c). The source proteins were also enriched for basement membrane compartment in the background of total XPT repertoire, suggesting association with extracellular matrixes (Figure 5d). This compartment could explain why these peptides are specifically presented in the context of cross-presentation, as basement membrane associated proteins are outside of plasma membrane and thus their access to the endogenous antigen processing machinery is limited. Source proteins of XPT-only antigens did not significantly overlap with the source proteins of tumor endogenous antigens (43.7%) or with the source proteins of immune cell endogenous antigens (21.7%); indeed, the majority of source proteins were unique to the XPT-only subset (Figure 5e). XPT-only peptides were largely derived from a diverse set of source proteins, with only 5% of source proteins containing multiple XPT epitopes (Figure 5f).

**Figure 5.**
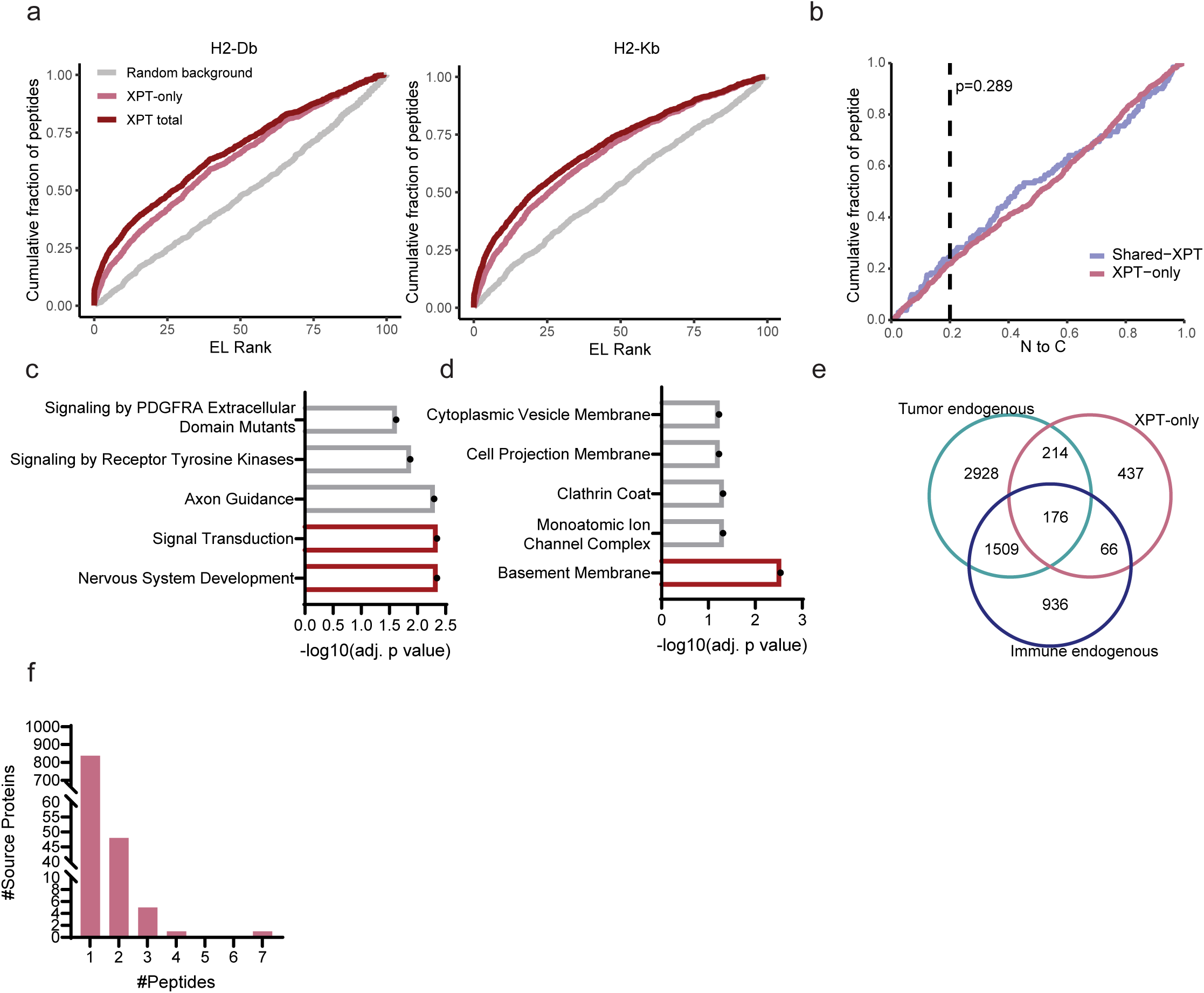
XPT-only antigens enrich for basement membrane associated proteins. a). Cumulative distribution function plot of netMHCPan predicted EL rank of total XPT peptides (red), XPT-only peptides (pink), and random peptides sampled from the proteome (grey) binding to H2-Kb (left) or H2-Db (right). b). Cumulative distribution function plot of relative position of XPT-only peptides (pink) and XPT-shared peptides (purple) within the source protein. Position was calculated using the position of the starting amino acid of the peptide. 0 represents N terminus and 1 represents C terminus. Fisher’s exact test was used to compare difference in ratios of XPT-tumor and shared -XPT peptides starting <20% from the N-terminus. c). GO biological process and d) GO cellular compartment enrichment analysis of source proteins from XPT-only antigens. Total XPT antigens associated source protein repertoire was used as enrichment background. P-value was adjusted by Benjamini-Hochberg method. Significant pathways (p<0.05) were colored in red. e). Venn diagram of XPT-only peptide, total immune endogenous peptide, and tumor endogenous peptide associated source proteins. f). Distribution of source proteins by number of XPT-only peptides contained.

### XPT-tumor peptides elicit strong immune response and delay tumor growth

Given the distinct characteristics of XPT antigens in different categories, we wanted to assess which of the XPT categories might be most effective at driving an anti-tumor immune response. To this end, we designed mRNA therapies encoding selected subsets of XPT antigens. Mice bearing GL261 subcutaneous tumors were administered 2 doses of each mRNA vaccine encoding the selected antigens or not treated (n=5), and ELISpot and flow cytometry was used to evaluate the immune response (Figure 6a). We initially compared XPT-only peptides to tumor endogenous peptides. Because XPT-only peptides are not endogenously presented on tumor cells, we reasoned that XPT-only peptides are unlikely to contribute to an effective anti-tumor response, as T cells primed against these antigens cannot target tumor cells that do not express these antigens. On the other hand, tumor endogenous peptides might enable a strong immune response, as these peptides are not typically presented on APCs and thus could represent novel antigens. To test this hypothesis, we selected 6 source proteins from which we had identified at least one XPT-only peptide and one tumor endogenous peptide (Supplementary Table 4). Using targeted PRM analysis, we first validated that the selected XPT-only targets were not presented endogenously on tumor cells or APCs, and that the tumor endogenous peptides were restricted to tumor cell presentation (Supplementary Figure 7). Mice were then vaccinated with an mRNA vaccine encoding either the 6 XPT-only peptides, or the corresponding 6 tumor endogenous peptides. A significantly weaker immune response was observed in mice vaccinated with the XPT-only mRNA vaccine compared to the tumor endogenous antigen mRNA vaccine (Figure 6b). However, the difference in antigen specific immune response did not result in any tumor growth rate difference (Supplementary Figure 8a). There was no significant difference in immunogenicity among individual peptides from either XPT-only antigens or tumor endogenous antigens (Supplementary Figure 8b). Both groups of antigens resulted in elevated levels of intratumoral CD8 effector T cells and decreased levels of PD1+ exhausted CD8 T cells compared to untreated mice, but no difference was observed between the two groups (Supplementary Figure 8c).

**Figure 6.**
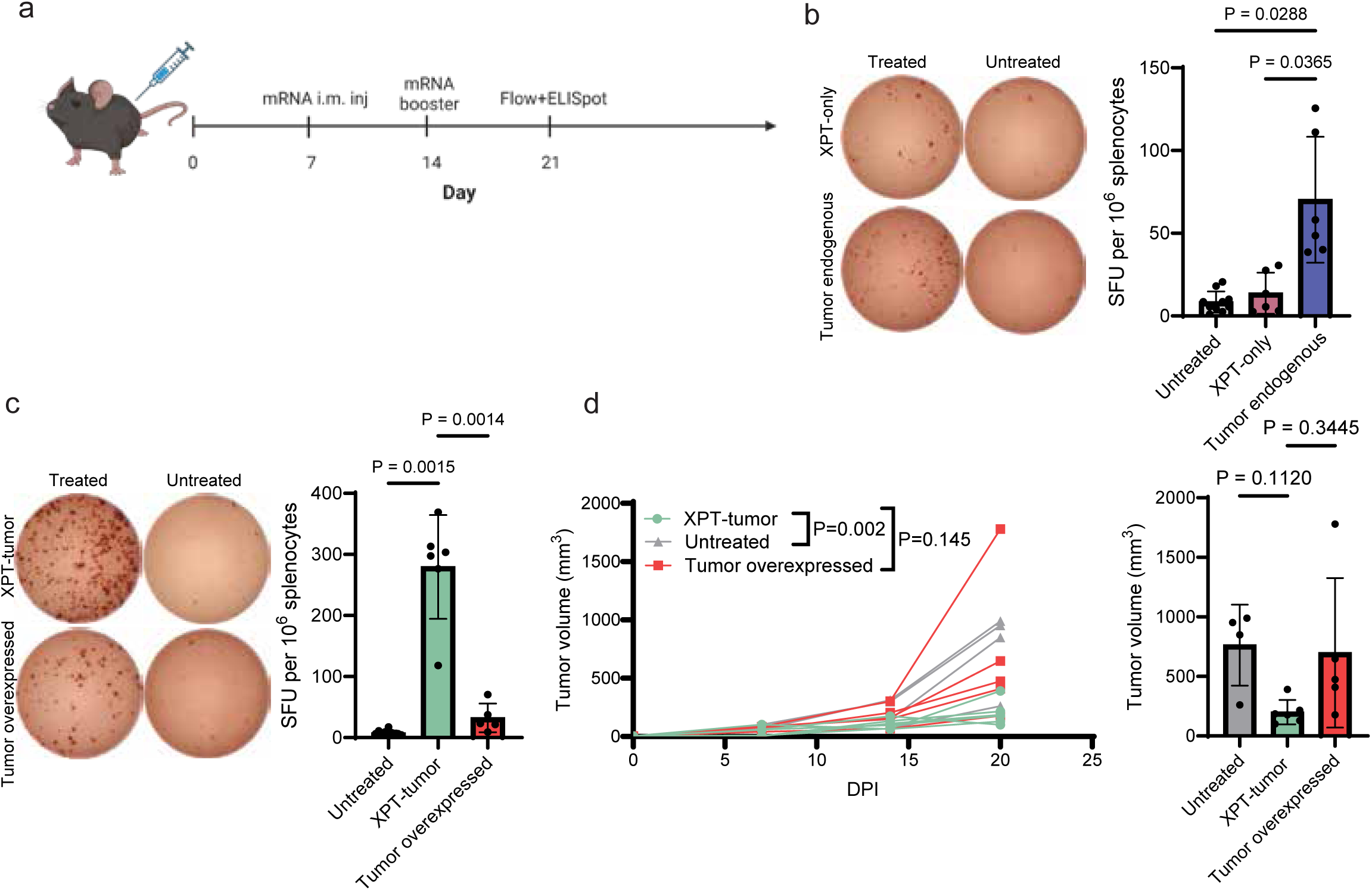
XPT-tumor peptides elicit strong immune response and delay tumor growth. a) Schematic of XPT antigen encoding mRNA vaccination. Mice bearing subcutaneous GBM tumors were injected with two doses of mRNA at D7 and D14 post tumor inoculation. b) (left) ELISPOT of splenocytes from XPT-only antigen treated, tumor endogenous antigen treated, and untreated mice stimulated with different groups of antigens. Representative image was selected from each treatment group. (right) Quantification of spots from each group of mice (n≥5). Significant difference among groups was determined by Brown-Frosythe and Welch ANOVA tests followed by multiple comparisons. p value was adjusted by Benjamini-Hochberg method. c) (left) ELISPOT of splenocytes from XPT-tumor antigen treated, tumor overexpressed antigen treated, and untreated mice stimulated with different groups of antigens. Representative image was selected from each treatment group. (right) Quantification of spots from each group of mice (n≥5). Significant difference among groups was determined by Brown-Frosythe and Welch ANOVA tests followed by multiple comparisons. p value was adjusted by Benjamini-Hochberg method. d) (left) Tumor growth curve comparison of XPT-tumor antigen treated, tumor overexpressed antigen treated, and untreated mice. Tumor growth was fitted using mixed-effect modeling, and the significance of difference in slopes among groups was determined by two-way ANOVA followed by multiple comparisons. p value was adjusted by Holm-Bonferroni method. (right) Tumor volume comparison at 20 DPI. Significant difference among groups was determined by Brown-Frosythe and Welch ANOVA tests followed by multiple comparisons. p value was adjusted by Benjamini-Hochberg method.

We next wanted to determine whether XPT-tumor antigens or other tumor associated antigens would be more immunogenic, and whether either of these could drive an immune response that limited tumor growth. With the goal of selecting more potent tumor-associated antigens, we selected 6 highly presented tumor associated antigens derived from genes overexpressed in human GBM tumors with low expression in normal brain and other tissues (‘tumor-overexpressed’) (Supplementary Figure 9a). For the XPT-tumor peptides, we selected 6 peptides derived from source proteins involved in tumor-associated processes (e.g., cell cycle or proliferation), or source proteins previously implicated in GBM tumors. XPT-tumor antigens and tumor-overexpressed endogenous antigens were made into mRNA vaccines and separately administered to mice bearing subcutaneous GL261 tumors (Figure 6a, Supplementary Table 5). The mRNA vaccine comprised of XPT-tumor antigens generated a significantly stronger immune response compared to the vaccine expressing tumor overexpressed antigens (Figure 6c). Stimulation of splenocytes with individual peptides from each group revealed that one peptide in particular from the XPT-tumor antigen group, SGLVNHVPL, was immunodominant (Supplementary Figure 9b). Aimp1, the source protein for this peptide, has been shown to be upregulated in high grade gliomas and negatively affects response to anti-angiogenic therapy^37^. Other XPT-tumor peptides elicited similar immune responses compared to peptides in the tumor overexpressed antigen group (Supplementary Figure 9c), suggesting that immunodominance is more of a feature of the Aimp1-derived peptide than a general feature of XPT-tumor peptides. It is worth noting that the immune response generated did not correlate with MHC-I binding affinity of the peptides (Supplementary Figure 9d). We also observed delay in tumor growth when mice were vaccinated with the XPT-tumor antigens, with average tumors 76.8% and 74.6% lower in volume compared to untreated mice and mice treated with tumor overexpressed antigen group 6 days after the second dose of mRNA (Figure 6d). However, we did not observe any difference in the effector or exhausted CD8 T cell populations between XPT-tumor antigen group and tumor overexpressed antigen group (Supplementary Figure 9e). All mRNA vaccinated groups showed higher percentage of effector CD8 T cells and lower percentage of exhausted T cells compared to the untreated group. Together, these results suggest that cross-presented antigens could elicit stronger immune response compared to tumor associated antigens that are not cross-presented, making them improved immunotherapy targets.

## Discussion

Cross-presentation in the tumor microenvironment has been extensively studied in recent years and has been shown to mediate anti-tumor response by priming antigen specific CD8+ T cells. However, MHC-I peptide level mapping in cross-presentation remains poorly characterized, but this information is critical for designing improved immunotherapies with targets that can be efficiently cross-presented by APCs. To generate this critical information, here we analyzed the XPT antigen repertoire in detail by profiling XPT GBM tumor antigens on BMDMs, BMDC, and splenic DCs.

In total, we identified over a thousand XPT peptides; these peptides distinctly differ from peptides endogenously presented by APCs or tumor cells. Only 15% of peptides in the XPT repertoire were shared with APC endogenous antigen presentation; this ratio of overlap was similar when compared against GL261 or CT2A endogenous antigen presentation. Additionally, 67.3% and 67.8% source proteins represented by the XPT antigens were different from the source proteins represented by APC or tumor endogenous antigens, respectively. XPT antigen source proteins were also enriched for biological processes different from those of either the APC or tumor endogenous peptide repertoires, highlighting transcriptional regulation associated pathways. Such differences in cross-presented and endogenously presented antigens has been reported by other studies. For instance, early studies in viral infection demonstrated variance in cross-priming and direct-priming efficiency among viral proteins, while cross-priming efficiency dictated their immunodominance hierarchy in CD8+ T cell responses, emphasizing that XPT fundamentally reshapes the epitope landscape visible to CD8 T cells^38,39^. Fessenden et al. also reported a distinct XPT repertoire using a SILAC-based melanoma–DC co-culture system^16^. The observations together support cross-presentation being a selective filtering process that yields a distinct antigen repertoire, instead of a diluted pool of the tumor immunopeptidome. The distinction between cross-presentation and tumor endogenous repertoire highlights the importance of characterizing antigen presenting cells when selecting immunotherapy targets rather than solely focusing on tumor presented antigens. Comparing the XPT repertoire with endogenous tumor and APC peptides, we categorized the XPT antigens into 3 categories: XPT-XPT-shared endogenous peptides that were also presented on tumor cells and APCs, XPT-only peptides that were not presented endogenously on either cell types, and XPT-tumor antigens that were specific to tumor endogenous presentation.

Consistent with other cross-presentation studies, we observed that the presentation of XPT peptides was lower compared to endogenous peptides presented on APCs, potentially due to a lower amount of tumor-derived source proteins relative to endogenous source proteins. To better understand the relationship between cross-presentation and endogenous presentation, we quantified copies/cell of XPT and endogenous peptide presentation for a selected subset of XPT-shared-shared peptides. This quantification revealed strong correlation between presentation of XPT-shared antigens and APC endogenous peptides, but not between XPT-shared antigens and tumor endogenous peptides, suggesting that XPT-shared antigens and APC endogenous peptides may be processed similarly in APCs. Additionally, XPT-shared antigens enriched more significantly for H2-Kb and Db motifs compared to total XPT peptides, suggesting pathway convergence on TAP preferences and tapasin editing with the canonical endogenous antigen processing pathways^40^. These findings support the cytosolic cross-presentation pathway model, where antigens are released from phagocytic compartments to the cytoplasm and further processed into short peptides by the proteasome^41^. These data also imply that endogenous peptides that are abundantly presented on APCs may be good candidates for efficient cross-presentation.

In the analysis of XPT-tumor peptides, we highlighted that there was little overlap between tumor endogenous antigens and XPT repertoire, and that cross-presentation of tumor endogenous antigens is less favored compared to shared antigens with APCs. This bias could result from differential antigen processing machinery during cross-presentation and tumor endogenous presentation. Inefficiency in cross-presenting tumor endogenous antigens may also play a role in defective T cell priming and activation in many solid tumors. Compared to APC endogenous antigens, XPT-tumor antigens more closely resemble tumor endogenous antigens in biophysical properties, displaying higher net charge, hydrophobicity, and isoelectric points. Additionally, XPT-tumor peptides had lower MHC-I binding affinity and were enriched at source protein N terminus compared to XPT-shared antigens. Among the peptides with low MHC-I binding affinity, we found over 10 XPT-tumor peptides from cytoskeletal protein vimentin, which supports a previous finding that cytoskeletal proteins are favored during cross-presentation through DNGR-1 detection of filamentous actin^42,43^. Although vimentin-derived peptides were predicted to bind poorly to MHC-I complex, we reasoned that such targets are unlikely to result from non-specific binding to the beads because other highly abundant cytoskeletal protein derived peptides were not identified in the XPT-tumor repertoire. These observations suggest that XPT-tumor epitopes are less trimmed and potentially originated from tumor derived degraded protein repertoire known to be enriched for N-terminal protein segments^44^. The identified XPT-tumor antigens support the model of vacuolar pathway cross-presentation, where epitopes are generated by cathepsins to load on MHC-I molecules in the phagosome without motif selection by chaperones in the peptide loading complex.

Intriguingly, the majority of the XPT repertoire was comprised of ‘XPT-only’ peptides that were not presented endogenously on tumor cells or APCs. The presentation of these novel XPT-only peptides could potentially be explained by different antigen processing pathways used in cross-presentation, leading to a distinctly different antigen repertoire compared to self-presentation. Additionally, enrichment analysis revealed that these XPT-only peptides were enriched for basement membrane proteins, suggesting that their absence in tumor cell or APC endogenous antigen repertoires could result from limited accessibility to the source antigens. These data support the concept that cross-presentation functions as an antigen discovery and danger-sensing system by expanding the tumor antigen repertoire, but whether these novel XPT antigens serve as ‘decoy receptors’ or whether they are able to generate an effective CD8+ T cell response requires further exploration.

To explore the in vivo immunogenicity of the identified XPT antigens, we made mRNA vaccines encoding 6 peptides from XPT-only antigens, tumor-presented antigens derived from the same source proteins as the XPT-only antigens, XPT-tumor antigens, and tumor overexpressed antigens. After 2 doses of each mRNA vaccine, mice bearing subcutaneous GL261 tumors generated a weaker immune response against the XPT-only antigens compared to the tumor presented antigens derived from the same source proteins, suggesting that the XPT-only antigens were unable to induce an effective CD8+ T cell response. Vaccination with mRNA comprised of XPT-tumor antigens elicited a strong antigen specific T cell response and delayed tumor growth compared to the mRNA comprised of tumor overexpressed antigens, suggesting that although the XPT-tumor antigens are rare, they could serve as potent immunotherapy targets. Interestingly, one epitope from the XPT-tumor set was highly immunogenic and dominated the antigen specific T cell response. This immunodominance was not correlated with MHC-I binding affinity, suggesting that immunogenicity is a complex characteristic determined by multiple factors including TCR activation, antigen processing efficiency and existing T cell repertoires of the host ^45,46^. Vaccination increased effector tumor infiltration of CD8 T cells and resulted in a lower percentage of exhausted CD8 T cells independent of epitope sequences, suggesting that mRNA is an effective onco-therapeutic modality. The similar percentage of CD8 T cell infiltration and exhaustion in mice vaccinated with XPT-tumor antigens and tumor overexpressed antigens suggests that further analysis is required to understand the mechanism of delayed tumor growth in the context of mRNA vaccination with XPT-tumor antigens.

## Conclusion

In this work we profiled the MHC-I immunopeptidome of cross-presented GBM antigens, providing, to our knowledge, the most comprehensive characterization of cross-presented MHC-I ligands in glioma. By combining heavy-isotope labeling with co-culture models of tumor-to-APC antigen transfer, and immunopeptidomic analysis, we defined over 1000 cross-presented GBM peptides on BMDCs, BMDMs, and splenic DCs after uptake of tumor material. We demonstrated that the cross-presented immunopeptidome cannot be inferred from endogenous presentation of tumor cells or APC alone, and could vary by antigen source, APC subtypes, and possibly tissue context. Importantly, we observed a striking bias against epitopes unique to tumor cell endogenous presentation during the cross-presentation process, highlighting the impact of APC imposed restrictions from antigen routing and processing on the XPT peptide repertoire. Our data suggest that subsets of the XPT repertoire are processed by the canonical APC antigen processing pathway, while others bypass this route and instead are processed through the vacuolar pathway. Therapeutically, our data suggest that antigens that are cross-presented on APC and endogenously presented on tumor cells, although rare in the context of all XPT peptides, may be the most immunogenic and may represent improved immunotherapy targets compared to tumor associated antigens. Together, our data fill a critical gap in the tumor immunology field and establish a novel antigen atlas for future studies of DC-based vaccines, myeloid-targeted immunotherapies, and antigen prediction algorithms.

## Methods

### Animals

C57BL/6 female mice were purchased from Jackson Laboratory (RRID:IMSR_JAX:000664). Mice 6-12 weeks old were acclimated for 3 days before any experiments. All animal procedures were approved by the Committee on Animal Care at MIT.

### Cell culture

GL261 cells (RRID:CVCL_Y003) and CT2A cells (RRID:CVCL_ZJ44) were purchased from ATCC in 2020. Cells were authenticated using Short Tandem Repeat (STR) profiling, and no re-authentication was performed after purchase. H2 null GL261 and CT2A cells were engineered using CRISPR knockout according to previous report^21^. The cells were grown in DMEM/F12 medium with 10% FBS at 37C, 5% CO2, and kept under 15 passages for all experiments. Mycoplasma was monitored using MycoAlert® Mycoplasma Detection Kit (Lonza) every 6 months.

To culture primary murine immune cells, bone marrow cells were harvested from femurs of C57BL/6 mouse according to published protocols^22^. After harvest, bone marrow cells were centrifuged at 300xg for 10 mins and red blood cells were lysed using RBC lysis buffer (Thermo Fisher) for 5 minutes at room temperature. Harvested bone marrow cells were frozen in 90% FBS with 10% DMSO (Sigma) and stored for up to 12 months. Bone marrow derived dendritic cells (BMDCs) were differentiated in complete RPMI media supplemented with 25mM HEPES (Gibco), 10% FBS, 1x β- mercaptoethanol (Gibco), 1x penicillin/streptomycin (Gibco), 100ng/ml Flt3L-Fc (BioXcell), and 5ng/ml GM-CSF (BioLegend #576306) for 6 days before co-culture experiments. Media was replenished during differentiation on Day 3. On Day 6, floating BMDCs were collected and matured in differentiation media supplemented with 20ng/mL DMXAA (Invivogen # tlrl-dmx) for 24hrs. Adherent BMDMs were replenished with differentiation media and cultured for another 24hrs.

### SILAC labeling of tumor cells

Stable isotope labeling of amino acids in tumor cell culture (SILAC) was done with a custom culture medium composition (Supplementary Table 1). DMEM without amino acids (US Biological) was supplemented with essential light amino acids, and heavy isotope-labeled amino acids tyrosine (^13^C_9_ or ^13^C_9_, ^15^N in some discovery analysis), asparagine (^13^C_4_, ^15^N_2_), phenylalanine (^13^C_9_, ^15^N) (Sigma Aldrich). The custom media was also supplemented with 10% FBS, 1% penicillin/streptomycin (Gibco), D-glucose (4.5g/L), NaHCO_3_ (3.7g/L) and 1 mM HEPES (Gibco), according to Corning DMEM media recipe. The media was used to culture GL261 or CT2A tumor cells for 72 hrs after tumor cells have been cultured for 1 passage in regular complete DMEM. Labeling was timed with primary immune cell differentiation so that both labeled tumor cells and the immune cells were used fresh for co-culture. Heavy isotope labeled-amino acid incorporation was assessed by global protein expression profiling of an aliquot of the labeled tumor cells.

### Co-culture with tumor cells and processing for immunopeptidomics analysis

For each replicate, 20e6 bone marrow cells were differentiated and matured to yield approximately 10e6 BMDMs and BMDCs. On Day 7, 10e6 SILAC labeled H2 null CT2A or GL261 tumor cells were added to differentiated macrophages or DCs to reach approximately 1:1 tumor cell: antigen presenting cell (APC) ratio. Labeled tumor cells were lifted using trypsin and washed in PBS 2x to avoid heavy isotope labeled amino acid carry over. BMDCs were collected, washed with PBS, and re-plated in a new plate with the labeled tumor cells. Adherent BMDMs were washed with PBS before labeled tumor cells were added. Plates were swirled to ensure complete cell mixing. Co-culture was harvest after 18hrs. To harvest the co-culture, floating BMDCs were collected by spinning down the media, and the rest of tumor cells and loosely attached BMDCs were lifted with trypsin at 37. For BMDM co-culture, cells were lifted with trypsin at 37, and then separately with 10mM EDTA since 10mM EDTA is toxic to CT2A cells. The tumor cell and APC mixture was spun down, washed with PBS, and frozen at −80 until subjected to immunopeptidomics analysis.

### Peptide synthesis

Heavy amino acid labeled peptides, or light peptides were generated at the Biopolymers & Proteomics core facility at MIT, or at Biosynth International, Inc. Synthesized peptides were cleaved using standard cleavage cocktail and purified to >95% using HPLC. Molecular weight of the peptide was confirmed using MALDI mass spectrometer (Bruker microflex). Heavy isotope labeled amino acids used for synthesis were purchased from Cambridge Isotope Laboratories, Inc.

### Generation of recombinant heavy isotope-labeled peptide MHCs (hipMHCs)

Heavy amino acid labeled peptides and positive control peptides (provided by manufacturer) were separately loaded on recombinant mouse Kb/Db easYmers® (Immunaware), according to the manufacturer’s protocol. The concentration of stable H2-Kb and H2-Dbcomplexes post loading was quantified using an adapted protocol from Flex-T HLA class I ELISA assay (Biolegend). Briefly, positive control peptide and heavy peptide loaded complexes in serial dilution were incubated in streptavidin coated plates. Stable complex was detected using HRP conjugated antibody against β2m protein (Biolegend #280303) and quantified using a Tecan plate reader Infinite 200 with Tecan icontrol version 2.0.0.0. after adding substrate solution. Generated hipMHCs were frozen in aliquots at −20 until use (less than 4 months).

### Peptide MHC (pMHC) isolation

Cell pellets of cross-presentation co-cultures or APC or tumor cell monocultures were resuspended in 1 mL MHC lysis buffer made of 20 mM Tris-HCl pH 8.0, 150 mM NaCl, 0.2 mM PMSF (Sigma), 1% CHAPS (Sigma), and 1x HALT Protease/Phosphatase Inhibitor Cocktail (Thermo Scientific), followed by brief sonication on ice (3 x 10 second microtip sonicator pulses at 30% amplitude) to disrupt cell membranes. Lysate was cleared by centrifugation at 16,000 g for 15 minutes at 4°C. Peptides bound to the MHC were isolated from the lysates by immunoprecipitation (IP) and size exclusion filtration, as previously described^21,23,24^. For each sample 0.1 mg of anti-mouse H2-Kb antibody (Y-3 clone, InVivoMAb #BE0172, RRID:AB_10949300) and 0.05mg of anti-mouse H2-Db antibody (produced in house) were conjugated to 20 μL FastFlow Protein A Sepharose bead slurry (Cytiva). Beads were washed 1x with IP wash buffer (20 mM Tris-HCl pH 8.0, 150 mM NaCl), followed by addition of lysate and if applicable, 100fmol of each hipMHC, and incubated rotating overnight at 4 °C to immunoprecipitate pMHCs. Beads were washed with 1x TBS and 2x HPLC grade water, and pMHCs were eluted using 10% acetic acid for 20 minutes at room temperature (RT). Peptides were isolated from antibody and MHC molecules using a 10K molecule weight cutoff filter (PALL Life Science). Alternatively, for some analysis peptides were eluted from pMHC complex using 1% Trifluoroacetic Acid (TFA) for 20 mins at RT and cleaned up using C18 stage tips (Empore, CDS Analytical) according to the manufacturer’s protocol. Elution from either MWCO or C18 cleanup was lyophilized and stored at −80 °C until analysis.

### Sample processing for analysis of labeled tumor cell proteome

Bicinchoninic Acid Assay (Pierce™ # 23225) was used to determine the concentration of SILAC labeled GL261 or CT2A samples. 100µg protein from each sample was processed through S-Trap™ sample processing technology according to manufacturer’s protocol (Protifi). Briefly, lysate was diluted in SDS (5% final concentration), reduced using DTT (Sigma # D9779-25g) and alkylated using IAA (Sigma # I1149-5G). Protein was collected in micro S-Trap column and cleaned using wash buffer (100mM TEAB in methanol, pH=8.0). Digestion was performed with trypsin at 47 for 1hr. Peptides were collected from the column and lyophilized for analysis.

### Mass spectrometry data acquisition

Samples were analyzed using Orbitrap Exploris 480 or Orbitrap Astral mass spectrometer (Thermo Fisher Scientific). Orbitrap Exploris 480 was coupled to an UltiMate 3000 RSLC Nano LC system (Dionex), Nanospray Flex ion source (Thermo Scientific), and column oven heater (Sonation). Orbitrap Astral mass spectrometer was coupled to an Vanquish™ Neo UHPLC System (Thermo Fisher Scientific). The peptide MHC sample was resuspended in 5µl of 3% acetonitrile, 0.1% Formic Acid, and loaded onto a 10-12 cm analytical capillary chromatography column with an integrated electrospray tip (∼1 μm orifice), prepared in house (50 μm ID & 1.9 μM C18 beads, ReproSil-Pur) through WPS-3000 autosampler (Dionex) for Orbitrap Exploris 480 acquisition and Vanquish Neo integrated autosampler for Orbitrap Astral acquisition.

DDA analyses of unlabeled pMHC samples on Orbitrap Exploris 480: Peptides were eluted using a gradient with 6-30% buffer for 62 minutes, 30-45% for 12 minutes, 45-55% for 8 minutes, 55-97% for 2 minutes, and 97% to 3% for 1 minute. Standard mass spectrometry parameters were as follows: spray voltage, 2.5 kV; no sheath or auxiliary gas flow; heated capillary temperature, 280 °C. Full scan mass spectra (300-1200 m/z, 60,000 resolution) were acquired in the orbitrap analyzer after accumulation of 3e6 ions (normalized AGC target 300%). For every full scan, up to 3 sec cycle time of ions were subsequently isolated if the ions reached a minimum intensity threshold of 5e3, were within precursor range 350-1200 m/z, and had charge state of 2-4. For MS2 acquisition, ions were collected with an isolation window of 0.4 m/z, 120,000 resolution, maximum injection time of 247 ms, normalized AGC target = 1000%, and fragmented by higher energy collisional dissociation (HCD) with a collision energy (CE): 30%. Ions were excluded for 30s after being acquired 2 times within 20s.

DDA analyses of unlabeled pMHC samples on Orbitrap Astral: Peptides were eluted using a gradient with 6-30% buffer for 66 minutes, 30-35% for 7 minutes, 35-55% for 5 minutes, 55-97% for 5 minutes, and 97% to 3% for 1 minute. Standard mass spectrometry parameters were as follows: spray voltage, 2.5 kV; FAIMS at standard with resolution with 3.5 L/min carrier gas flow; heated capillary temperature, 280 °C. Full scan mass spectra (300-1200 m/z, 60,000 resolution) were acquired in the orbitrap analyzer after accumulation of 3e6 ions (normalized AGC target 300%) or 100ms maximum injection time, at −40 and −60 FAIMS CV and 45% RF lens. For every full scan, up to 20 scans (−40 FAIMS CV full scan) or 100 scans (−60 FAIMS CV full scan) of ions at were subsequently isolated if the ions had charge state of 2-4. For MS2 acquisition, ions were collected with an isolation window of 1.4 m/z, and analyzed in the Astral analyzer with maximum injection time of 50 ms, normalized AGC target = 50%, and fragmented by higher energy collisional dissociation (HCD) with a collision energy (CE): 34%. Ions were excluded for 15 s after being acquired 1 time with +/- 5 ppm mass tolerance. Dependent scan was performed in single charge state per precursor only.

DDA analyses of unlabeled proteome samples on Orbitrap Exploris 480: Peptides were eluted using a gradient with 6-19% buffer B for 38 minutes, 19-29% for 17 minutes, 29-41% for 9 minutes, 41-97% for 3 minutes, and 97% to 3% for 1 minute. Standard mass spectrometry parameters were as follows: spray voltage, 2.5 kV; no sheath or auxiliary gas flow; heated capillary temperature, 275 °C. Full scan mass spectra (380-2000 m/z, 60,000 resolution) were detected in the orbitrap analyzer after accumulation of 3e6 ions (normalized AGC target of 300%) or 25 ms. For every full scan, up to 3sec cycle time of ions were subsequently isolated if the ions reached a minimum intensity threshold of 1e4 and has charge state of 2-6. For MS2 acquisition, ions were collected with an isolation window of 0.4 m/z, 120,000 resolution, first mass at 100 m/z, maximum injection time of 247 ms, normalized AGC target = 100%, and fragmented by higher energy collisional dissociation (HCD) with a collision energy (CE): 30%. Ions were excluded for 30s after being acquired 2 times within 25s.

SureQuant Survey analyses: Peptides were eluted using a gradient with 5-25% buffer B (70% Acetonitrile, 0.1% formic acid) for 55 minutes, 25-45% for 22.5 minutes, 45-97% for 2.5 minutes, and 97% to 3% for 1 minute. Standard mass spectrometry parameters were as follows: spray voltage, 2.5 kV; no sheath or auxiliary gas flow; heated capillary temperature, 275 °C. The Mass spectrometer was operated in data dependent acquisition (DDA) mode with an inclusion list of the heavy trigger peptide at charge states 2-4. Full scan mass spectra (300-1500 m/z, 60,000 resolution) were detected in the orbitrap analyzer after accumulation of 3e6 ions (normalized AGC target of 300%) or 50 ms. For every full scan, up to 20 ions were subsequently isolated if the m/z was within +/- 5 ppm of targeted trigger peptide m/z. MS/MS spectra of targeted ions were collected with 120,000 resolution, 350-1500 scan range, a maximum injection time of 250ms, normalized AGC target = 1000%, and fragmented by higher energy collisional dissociation (HCD) with a collision energy (CE): 30%. Library of acquired spectra of each peptide at the charge state that generated the highest precursor intensity was created using Skyline software for SureQuant targeted analyses.

SureQuant targeted analyses: Peptide elution gradient for targeted analyses was the same as that of the survey analyses. The custom SureQuant acquisition template available in Thermo Orbitrap Exploris Series 2.0 was utilized for building this method. All the acquisition parameters for heavy labeled peptides were located within a distinct 5-node branch stemming from a full scan node. Each branch houses information of trigger peptides with the same heavy labeled amino acid at the same charge states, and the same amino acid SILAC labels corresponding to the XPT version of the peptides. In the full scan, the trigger peptide m/z and intensity thresholds are defined in the “Targeted Mass” filter node as 1% of the intensity from DDA survey run. Next, parameters for the low resolution, trigger peptide MS2 scan are specified, followed by the “Targeted Mass Trigger” filter node, which defines the 6 product ions used for pseudo-spectral matching (Supplementary Table 2). To connect each set of product ions within the targeted mass trigger node to a given precursor mass, group ID feature was used to specify the precursor m/z associated with each group of product ions. Below the trigger node, a few MS2 scans were acquired. First, MS2 scan of the endogenous peptide was acquired at the corresponding isolation offset (m/z). Additionally, MS2 scans of the XPT peptides with all SILAC labeled forms detected in previous DDA runs were acquired. The MS2 scans of the endogenous peptides and XPT peptides were defined in the scan parameters within each node. Standard mass spectrometry parameters for SureQuant acquisition are as follows: spray voltage: 2.5kV, no sheath or auxiliary gas flow, heated capillary temperature: 280°C. Full-scan mass spectra were collected with a scan range: 380-1200 m/z, AGC target value: 300% (3e6), maximum injection time: 50 ms, resolution: 120,000. Heavy peptides matching the m/z with 5 ppm tolerance and exceeding the intensity threshold defined on the inclusion list were isolated with isolation window 0.4 m/z and fragmented with 30% normalized collision energy by HCD with a scan range: 150-1200 m/z, maximum injection time: automatically determined from the resolution, AGC target value: 1000% (10e6), resolution: 60,000. A product ion trigger filter next performs pseudo-spectral matching, only triggering an MS2 event of the endogenous target peptide at the defined mass offset if n ≥ 4 product ions are detected from the defined list with 10 ppm mass tolerance. If triggered, the subsequent endogenous peptide and XPT peptides MS2 scans are initiated at the defined mass offsets. Scan parameters have the same collision energy, scan range, and AGC target as the heavy trigger peptide, but with a higher maximum injection time and resolution (max injection time: 247 ms, resolution, 120,000).

### SureQuant data analysis

Skyline software (RRID:SCR_014080) was used to quantify abundance of the standard and endogenous peptide. For each sample, the abundance of each target peptide (heavy, light, and XPT) was approximated using an average of the maximum intensity of top 3 product ions across the elution chromatogram. The absolute amount of endogenous and XPT peptide was calculated using the ratio of light:heavy abundance (heavy peptides were spiked in at 100fmol). To eliminate potential contamination of heavy peptide intensity in the isolation window of light or XPT peptides, ratio of baseline intensity of each product ion across the elution chromatogram at each isolation offset corresponding the endogenous and XPT peptides to the heavy product ion intensity was calculated from a SQ acquisition analyses with only the heavy peptides. The baseline ratio was subtracted from each endogenous and XPT peptides, and copies of endogenous peptide per cell were calculated using the absolute amount and input cell number in the IP. If multiple SILAC labeled versions of the XPT peptide existed, the copies/cell were summed to represented the total XPT level (Supplementary Table 3).

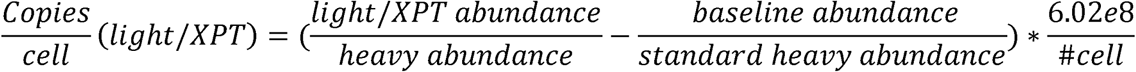

### Mass spectrometry data search

All mass spectra from DDA MS were analyzed with Proteome Discoverer (PD, version 3.0, RRID:SCR_014477). Co-culture samples acquired on Orbitrap Exploris 480 or Orbitrap Astral and monoculture samples acquired on Orbitrap Astral were searched using Mascot Server 3.0 against the Swiss-Prot mouse database (Identifier: Mus musculus, UniProt release 2024_06). Monoculture samples acquired on Orbitrap Exploris 480 were searched using SEQUEST against the Swiss-Prot mouse database (Identifier: Mus musculus [sp_canonical TaxID=10090_and_subtaxonomies], release 2024_06) and rescored using INFERYS. No enzyme was used, variable modifications included oxidized methionine for all analyses. In monoculture analysis with SEQUEST, static modification of Carbamidomethyl was used. Co-culture samples were searched with additional dynamic modifications of SILAC labels on tyrosine, phenylalanine, and asparagine. All analyses searched on SEQUEST were filtered with the following criteria: search engine rank = 1, length between 8 and 12 amino acids, q value ≤ 0.05. All analyses searched on Mascot were filtered with the following criteria: search engine rank = 1, length between 8 and 12 amino acids, Ion Score ≥ 15.

### modRNA production for mRNA vaccine

The mRNA vaccine sequences were designed by linear connection of 3 peptide sequences (Supplementary Table 4-5), where each peptide was separated by AAY linker to ensure proteasomal cleavage. Template DNA plasmids used in the production of modRNA were created using a commercially available Cloning Kit for mRNA Templates (Takara #6143) according to manufacturer’s instructions. Resultant plasmid DNA was linearized via endonuclease digestion and purified with PureLink PCR Purification columns (ThermoFisher #K310002) following manufacturer’s instructions. To synthesize RNA, 20μL in vitro transcription (IVT) reactions were performed using reagents from the HiScribe T7 High Yield RNA Synthesis Kit (NEB #E2040) and 1-2μg of linear DNA template (scaled as needed). Importantly, modified base N1-methylpseudouridine triphosphate (TriLink #N-1081) was added to the reaction mixture instead of canonical uridine triphosphate, and CleanCap Reagent AG (TriLink #N-7113) was utilized to co-transcriptionally add 5’ Cap-1 structures to synthesized RNA. The IVT product was purified using PureLink RNA Mini columns (ThermoFisher #12183018A) following manufacturer’s instructions. Quality of the resulting modRNA was assessed using UV-Vis spectrophotometry and gel electrophoresis.

### Lipid nanoparticle synthesis

9-Heptadecanyl 8-{(2-hydroxyethyl)[6-oxo-6-(undecyloxy)hexyl]amino}octanoate (SM-102) was purchased from BroadPharm (CAT#BP-25499); 1,2-distearoyl-sn-glycero-3-phosphocholine (DSPC; CAT#850365), Cholesterol (CAT#700100), 1,2-dimyristoyl-rac-glycero-3-methoxypolyethylene glycol-2000 (DMG-PEG2k; CAT#88015) were purchased from Avanti Polar Lipids. Citrate buffer (pH 3; CAT#J61391-AK) was purchased from Alfa Aesar. For dialysis, 20K MWCO Slide-A-Lyzer™ MINI Dialysis Device (ThermoFisher Scientific), and RNAse-free PBS (AM9625; ThermoFisher Scientific) were used. Lipid nanoparticles were synthesized using a microfluidic organic-aqueous precipitation method. The organic phase was prepared by mixing the lipids SM-102, DSPC, Cholesterol, and DMG-PEG2k in ethanol at a molar ratio of 50:10:38.5:1.5. The aqueous phase of RNA was prepared by diluting the RNA (stored in RNAse-free water) with 10 mM citrate buffer at pH 3.0. The two phases were prepared at an ethanol:aqueous volume ratio of 1:3, and at an N:P ratio of 5:1. Each phase was loaded into a syringe (BD), and locked onto the NxGen microfluidic cartridge for mixing using a NanoAssemblr Ignite instrument (Precision Nanosystems). The Ignite was set to operate with the following settings: volume ratio- 3:1; flow rate- 12 ml/min; waste volume- 0 mL. The resulting LNPs were dialyzed against 20mM Tris Acetate with 8% sucrose using 10K MWCO Slide-A-Lyzer™ MINI Dialysis casettes (ThermoFisher Scientific) at 25°C for 4 hours.

### Subcutaneous tumor inoculation, vaccination, and monitoring

GL261 tumor cells were cultured for 2 passages before inoculation. Confluent tumor cells were lifted using trypsin and washed with PBS for 2 times. The cells were resuspended to a density of 100e6 cells/ml in PBS and injected on the right flank of C57BL/6 mice at 100µl per mouse (10e6 per mouse). Mice were randomized into different groups with 5 mice per group post tumor injection by another lab member blinded from the injection process. No criteria were set for excluding or including particular animals. No data point was excluded. Animal cage locations or sequence of receiving treatment was randomized by the person executing the experiment to minimize confounders. mRNA vaccine (10µg/dose) was injected intramuscularly on the right hindlimbs at Day 7 and Day 14 post tumor inoculation. Tumor size was measured using digital caliper once per week to monitor tumor growth until Day 20. mice were euthanized Day 21 post tumor inoculation for flow cytometry and ELISpot characterizations.

### Tissue processing for ELISPOT, and flow cytometry

Tumors, spleens, and inguinal lymph nodes were resected and stored in RPMI 1640 media (Gibco) on ice before further processing. Spleens were mashed through 70µm filters to obtain a single-cell solution and red blood cells were lysed with RBC lysis buffer (ThermoFisher) for 5min at room temperature. Splenocytes were washed twice with RPMI before further processing. Tumors and inguinal lymph nodes were digested with 125 U/ml collagenase IV and DNase in complete RPMI media with 10mM HEPES at 37°C for 30 mins with pestle mashing in between. Digested tumors and lymph nodes samples were strained through 70µM strainer (Corning) and resuspended in FACS buffer (2% FBS[Thermo Fisher Scientific] and 2mM EDTA[Gibco]) in PBS (Corning) and RBC lysed for 5 mins at room temperature.

### Flow cytometry

For flow cytometry staining, cells from tumor and lymph nodes were plated in U bottom well paltes. Cells were blocked with TruStain FcX™ PLUS (anti-mouse CD16/32) Antibody (Biolgend #156603, RRID:AB_2783137) diluted in FACS buffer for 10min on ice to prevent non-specific binding. Together during blocking, cells were stained with LIVE/DEAD™ Fixable Near IR viability stain (Thermo #L34992). Cells were then washed with FACS buffer and stained for surface proteins using a cocktail of fluorophore-conjugated antibodies (Supplementary Table 6) resuspended in FACS buffer. Cells were fixed with Fix/Perm wash buffer (BD Biosciences) for 30 min on ice, washed 2x with FACS buffer, and stained with a cocktail of fluorophore-conjugated antibodies against intracellular targets. Finally, the cells were washed with FACS buffer 2x before acquisition on flow cytometer. Sample acquisition was performed on BD flow cytometers (FACS Symphony A3) and analyzed using FlowJo v10 (TreeStar). For tumor infiltrating lymphocyte analysis, cells were pre-gated on singlets, live, CD8+, TCRb+, and CD44 high.

### IFNγ Elispot assay

ELISpot plates (EMD Millipore) were coated overnight at 4°C with anti-IFNγ capture antibody (BD Biosciences, RRID: AB_2868944). Plates were washed and blocked with complete RPMI1640 media for 2h at room temperature. Splenocytes were plated in complete RPMI media at 1e6 cells/well with either 10µg/well mixture of vaccinated targets, 10µg/well single vaccinated peptide targets, negative control (complete media) or positive control (complete media supplemented with 100 ng/mL PMA (Sigma-Aldrich) and 1 mg/mL ionomycin (Sigma-Aldrich)). Plates were incubated overnight (18h) at 37°C and 5% CO2 and developed using a mouse IFNγ ELISpot kit (BD Biosciences # 551083), following manufacturer’s instructions. After drying (overnight at RT), spot counts were determined using an ImmunoSpot (CTL) Elispot reader.

### Statistical analysis

Analyses were performed using GraphPad Prism 10 (RRID:SCR_002798), python and R. Comparisons between groups were performed using parametric (student’s t test) or non-parametric tests (Mann-Whitney U test, Kruskal-Wallis test) after F-test, where suitability of the test was assessed based on data distribution using visual methods. Comparison across multiple groups were performed using Brown-Forsythe and Welch ANOVA tests. Correction for multiple comparisons was performed where applicable. False discovery rates were calculated using Benjamini-Hochberg method. P values < 0.05 were considered statistically significant (*p < 0.05; **p < 0.01; ***p < 0.001; ****p < 0.0001; ns = not significant). GibbsCluster 2.0 was used for motif analysis. NetMHCpan 4.1 was used to determine pMHC binding affinity.

## Data availability

The mass spectrometry proteomics data have been deposited to the ProteomeXchange Consortium via the PRIDE partner repository with the data set identifier PXD072705. Mass spectrometry data were processed using R (4.3.1). All other raw data generated in this study are available upon request from the corresponding author.

## Figure Legend

**Supplementary Figure 1.** a). Receptor-ligand interactions that suggest XPT identified among macrophage, DC, tumor, and T cell clusters. Interactions filtered by p<0.01. Color scale shows probability of communication. b). Schematic of labeling ratio calculation in SILAC labeled GL261 and CT2A cells. c). Sequence motif enrichment of endogenous GBM tumor peptides (left), and fraction of endogenous peptides on APCs that contain selected SIL amino acids Y, F or N. d). Fraction of CT2A and GL261 tryptic peptides in global protein expression profiling with full, partial or no SILAC labeling. e). Sequence motif enrichment of total XPT peptides identified from BMDM and BMDCs. f). Cumulative distribution function plot of netMHCPan predicted EL rank of total XPT peptides (red), total APC endogenous peptides (blue), total tumor endogenous peptides (green), and random peptides sampled from the proteome (grey) binding to H2-Kb (left) or H2-Db (right). Black line represents EL rank=2.g). Upset plot of XPT antigens from different APC cell types cross-presenting different GBM tumor cell types. h). Heatmap of top10 processes from GO biological process enrichment analysis of source proteins from XPT antigens, endogenous APC antigens, and endogenous tumor antigens. Total proteome was used as enrichment background. P-value was adjusted by Benjamini-Hochberg method. Color shows - log10 adjusted p value.

**Supplementary Figure 2.** a). Venn diagram of endogenous peptides profiled from BMDM, BMDC and splenic DC. b). Percentage of endogenous peptides containing Y, F or N from BMDM, BMDC, and splenic DCs. c). Percentage of XPT peptides in the total profiled peptides on cross-presenting BMDMs, BMDCs, and splenic DCs. d). Number of profiled XPT peptides from BMDMs, BMDCs, and splenic DCs. e). Venn diagram of XPT peptides and associated source proteins profiled from BMDM, BMDC and splenic DC. f). GO cellular compartment enrichment analysis of source proteins from splenic DC XPT antigens. Total proteome was used as enrichment background. P-value was adjusted by Benjamini-Hochberg method. Significant pathways (p<0.05) were colored in red.

**Supplementary Figure 3.** a). Correlation of precursor abundance rank of XPT-shared peptides in different MS analysis. b). Ranked precursor abundance of peptides in a cross-presentation analysis. Selected XPT antigens are shown in blue and corresponding endogenous antigens are shown in black.

**Supplementary Figure 4.** Mirror plots of cross-presented antigen and trigger peptide standards, and product ion chromatogram matches of trigger peptide standard, cross-presented antigens, and endogenous antigens.

**Supplementary Figure 5.** Bar plot of cross-presentation and endogenous presentation of selected XPT-shared antigens in copies/cell level.

**Supplementary Figure 6.** a). Venn diagram of XPT-tumor peptides from GL261 or CT2A co-culture. b). Detection of XPT-tumor peptides in tumor cells or APC endogenous peptide repertoire in PRM analysis. c). EL rank distribution of Vimentin derived XPT-tumor peptides binding to H2-Kb or H2-Db. d). Ranked number of endogenous tumor endogenous peptides associated with different source proteins. Source proteins Atp5pf, Psmc1, Psmc4, and Fau that had multiple XPT-tumor peptides were highlighted in red. e). Ranked precursor abundance of peptides in a GBM tumor cell global proteomic analysis. Atp5pf, Psmc1, Psmc4, and Fau are shown in black and the ones with multiple XPT-tumor peptides were shown in blue. f). GO cellular compartment enrichment analysis of source proteins from XPT-tumor (top) antigens and XPT-shared antigens (bottom). Total XPT antigens associated source protein repertoire was used as enrichment background. P-value was adjusted by Benjamini-Hochberg method. Significant pathways (p<0.05) were colored in red. g). Biophysical properties of XPT-tumor peptides compared to endogenous APC peptides or endogenous tumor endogenous peptides. Kruskal-Wallis Test was used to determine significant difference among the 3 groups. Wilcoxon test was used for pairwise comparison. ***: p<0.001, ****: p<0.0001

**Supplementary Figure 7.** Detection of XPT-only peptides and tumor endogenous peptides from corresponding source proteins in tumor cells or APC endogenous peptide repertoire in PRM analysis.

**Supplementary Figure 8.** a) Tumor growth curve comparison of XPT-only antigen treated, tumor endogenous antigen treated, and untreated mice. b). Quantification of spots from XPT-only antigen treated (left) and tumor-endogenous antigen treated (right) mice stimulated with individual antigens. c) Percentage of CD8+TCRb+ cells in the tumor infiltrating lymphocytes (left) and percentage of PD1+ exhausted T cells in the CD8 T cell population (right)

**Supplementary Figure 9** a) Log2 CPM distribution of GBM tumor overexpressed gene in normal tissues, primary, and recurrent GBM tumors. b) Quantification of spots from XPT-tumor antigen treated mice stimulated with individual antigens. c) Quantification of spots from XPT-tumor antigen treated (left, excluding Aimp1 peptide) and tumor overexpressed antigen treated (right) mice stimulated with individual antigens. d) Percentage of CD8+TCRb+ cells in the tumor infiltrating lymphocytes (left) and percentage of PD1+ exhausted T cells in the CD8 T cell population (right)

