## Supplemental Figures for "Analysis of tumor-derived and cross-presented peptide antigens defines improved immunotherapeutic strategies"

### Supplement 1

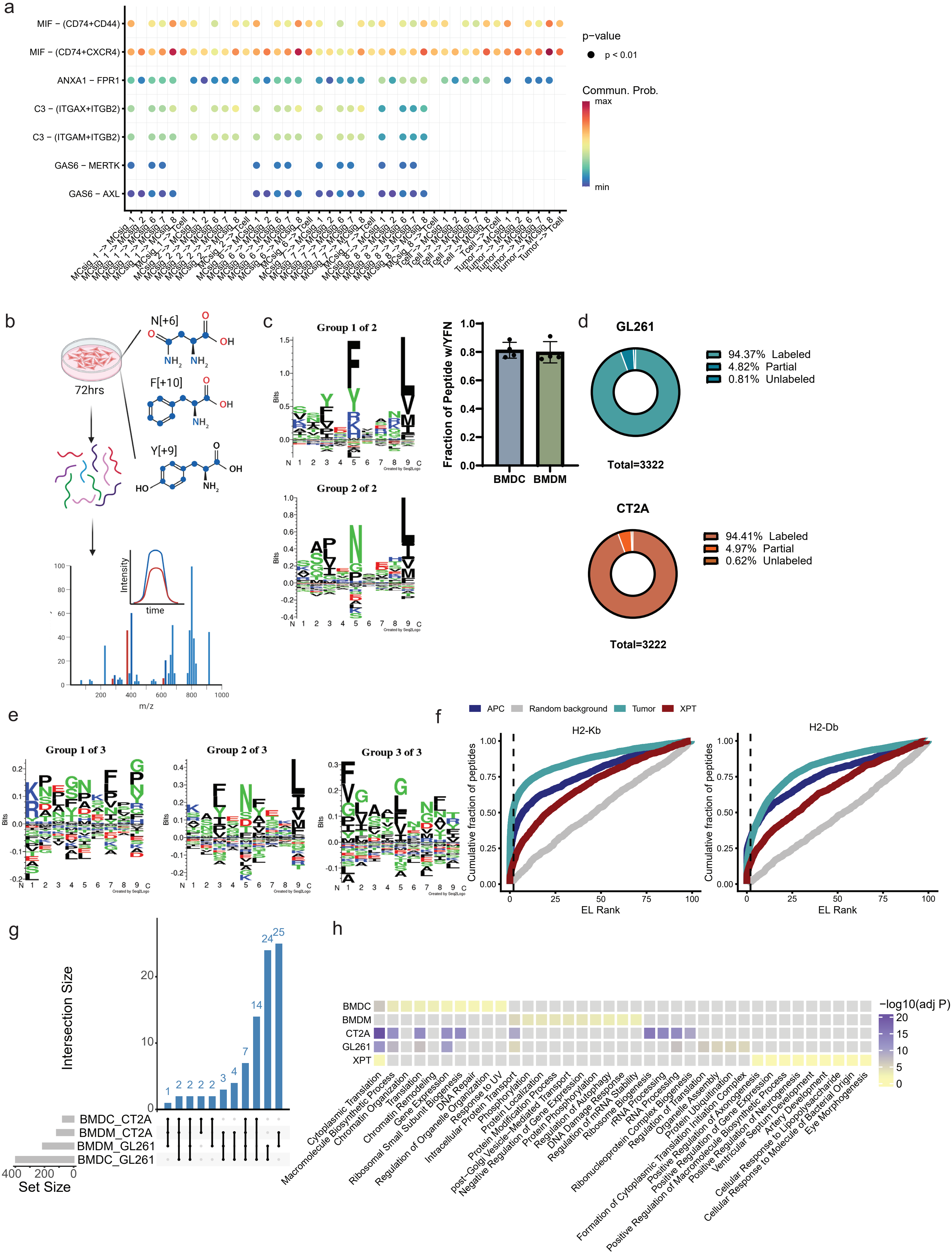

### Supplement 2

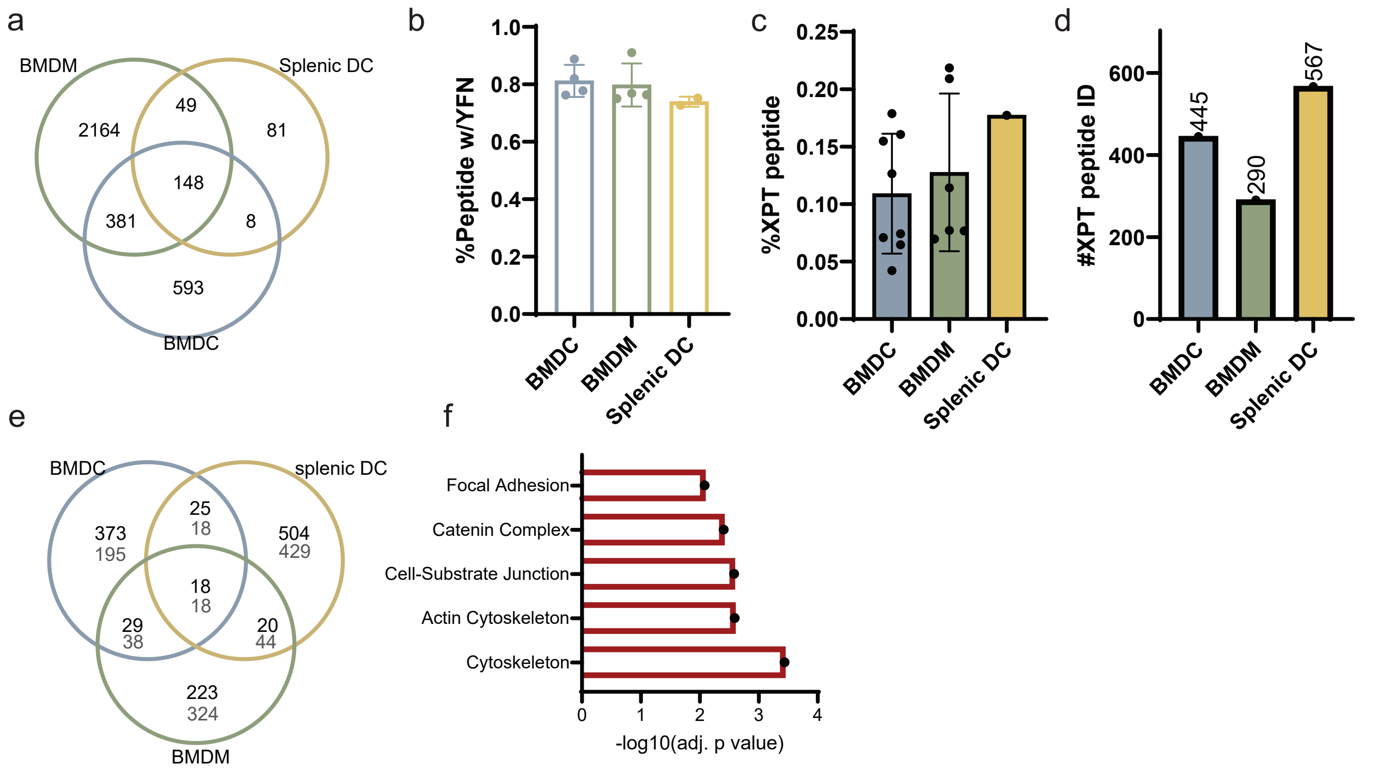

### Supplement 3

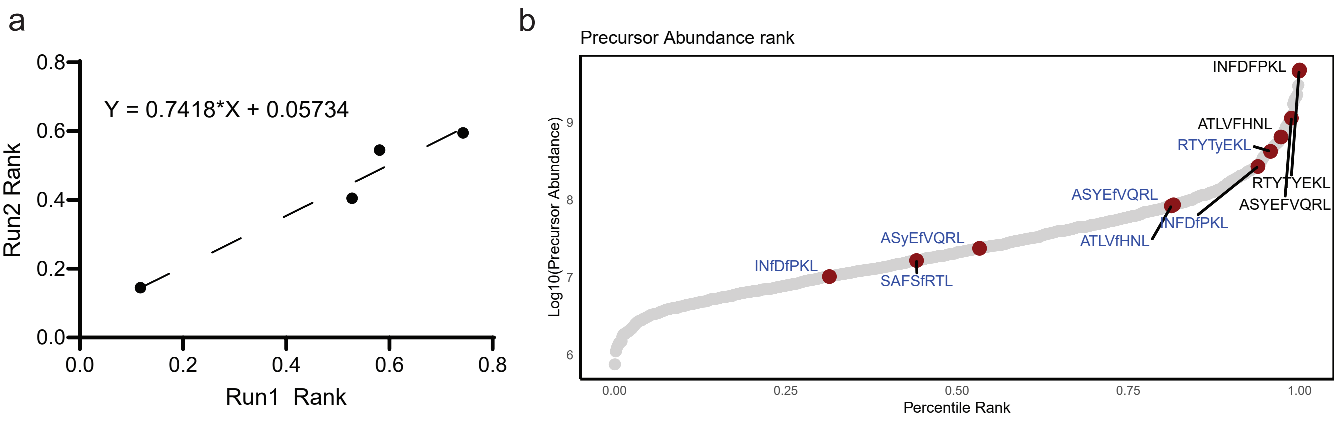

### Supplement 4

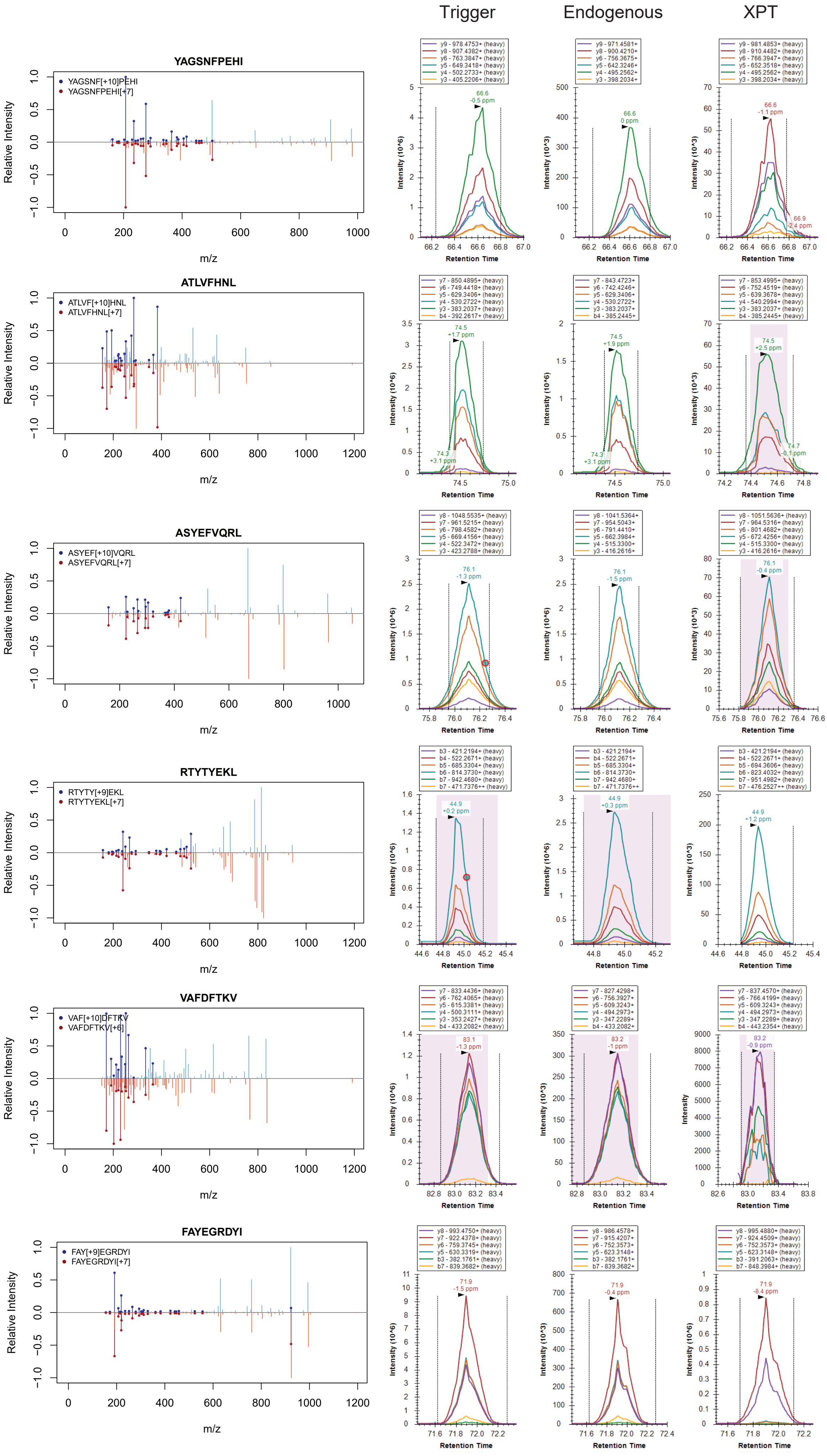

### Supplement 4

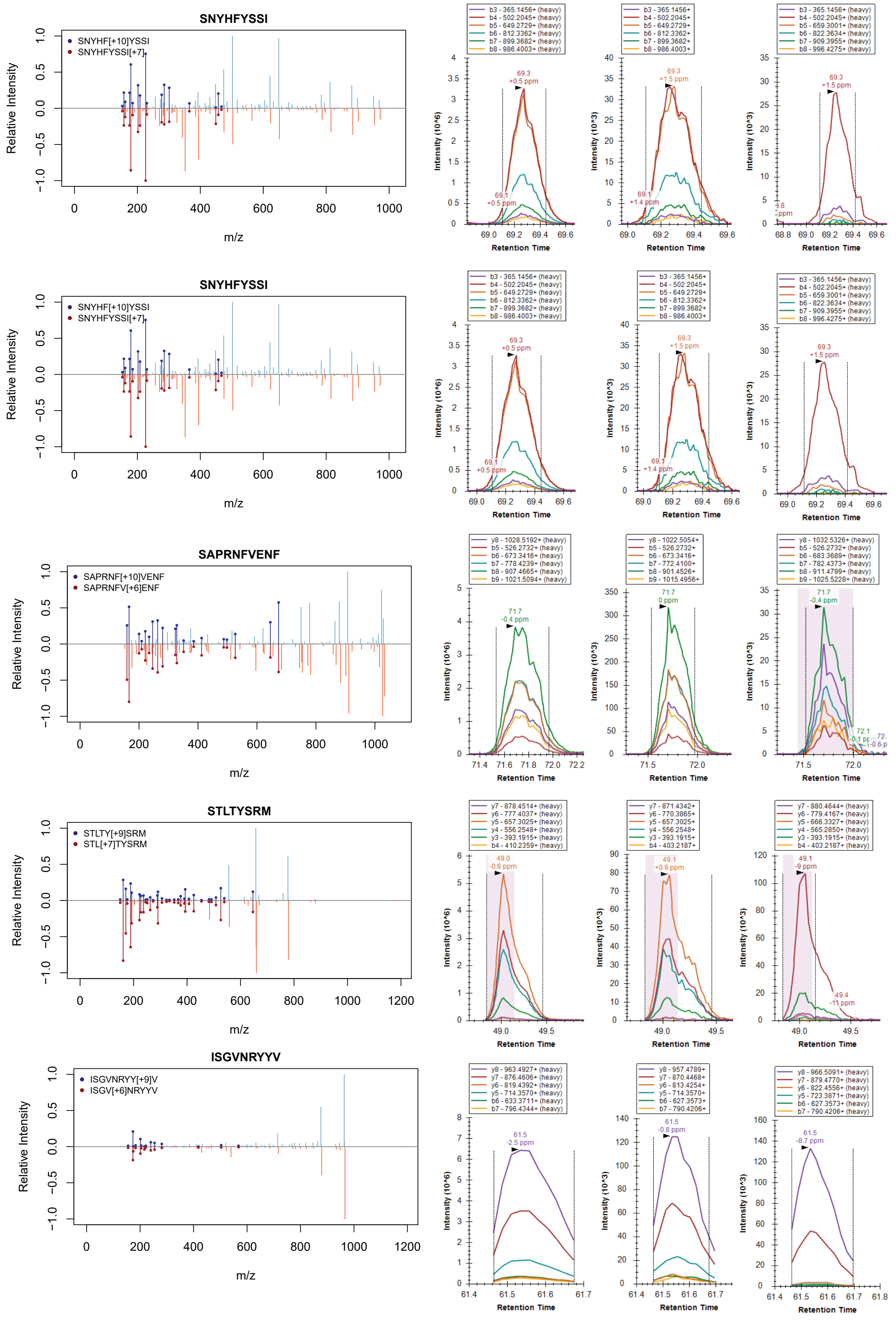

### Supplement 5

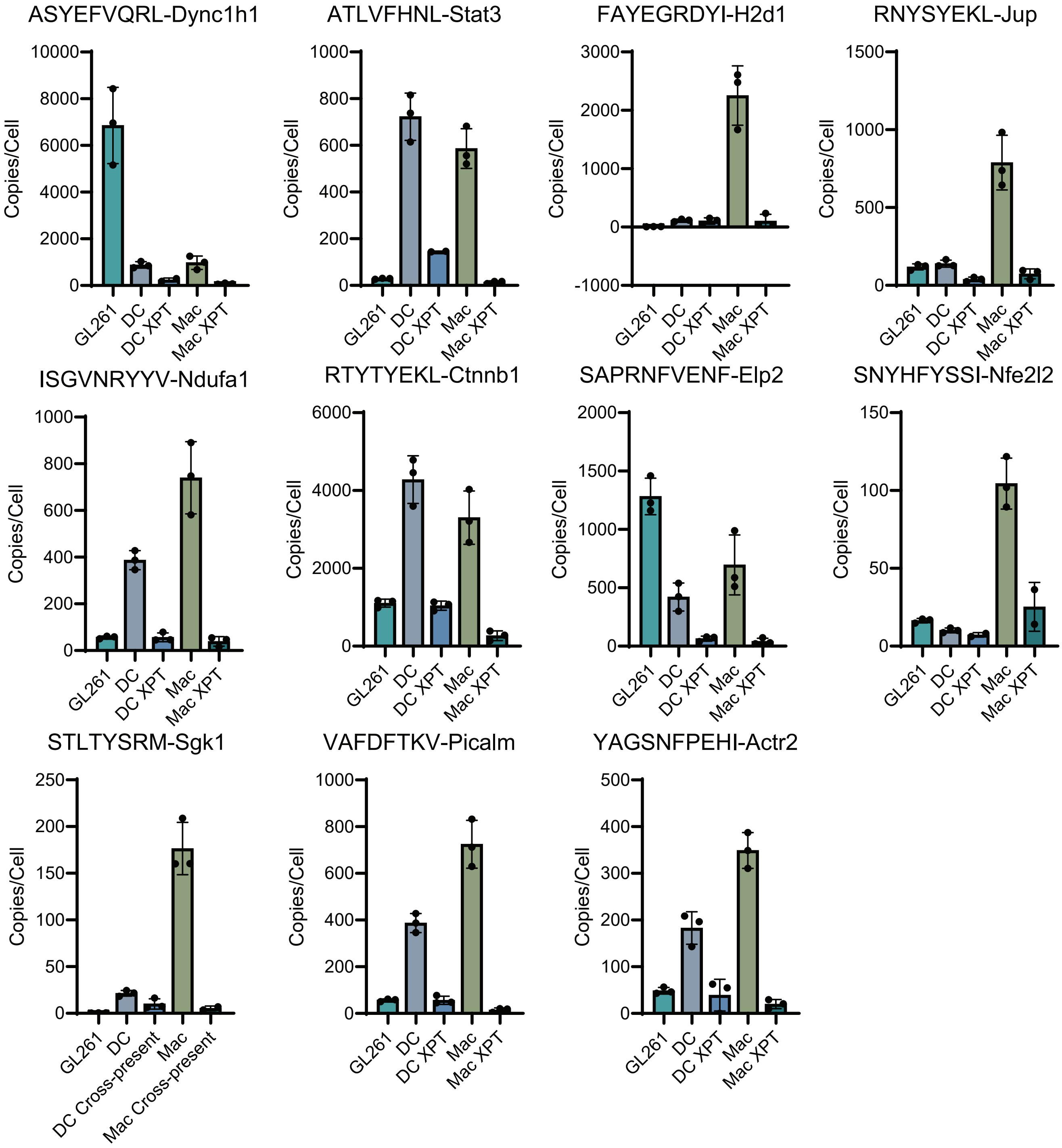

### Supplement 6

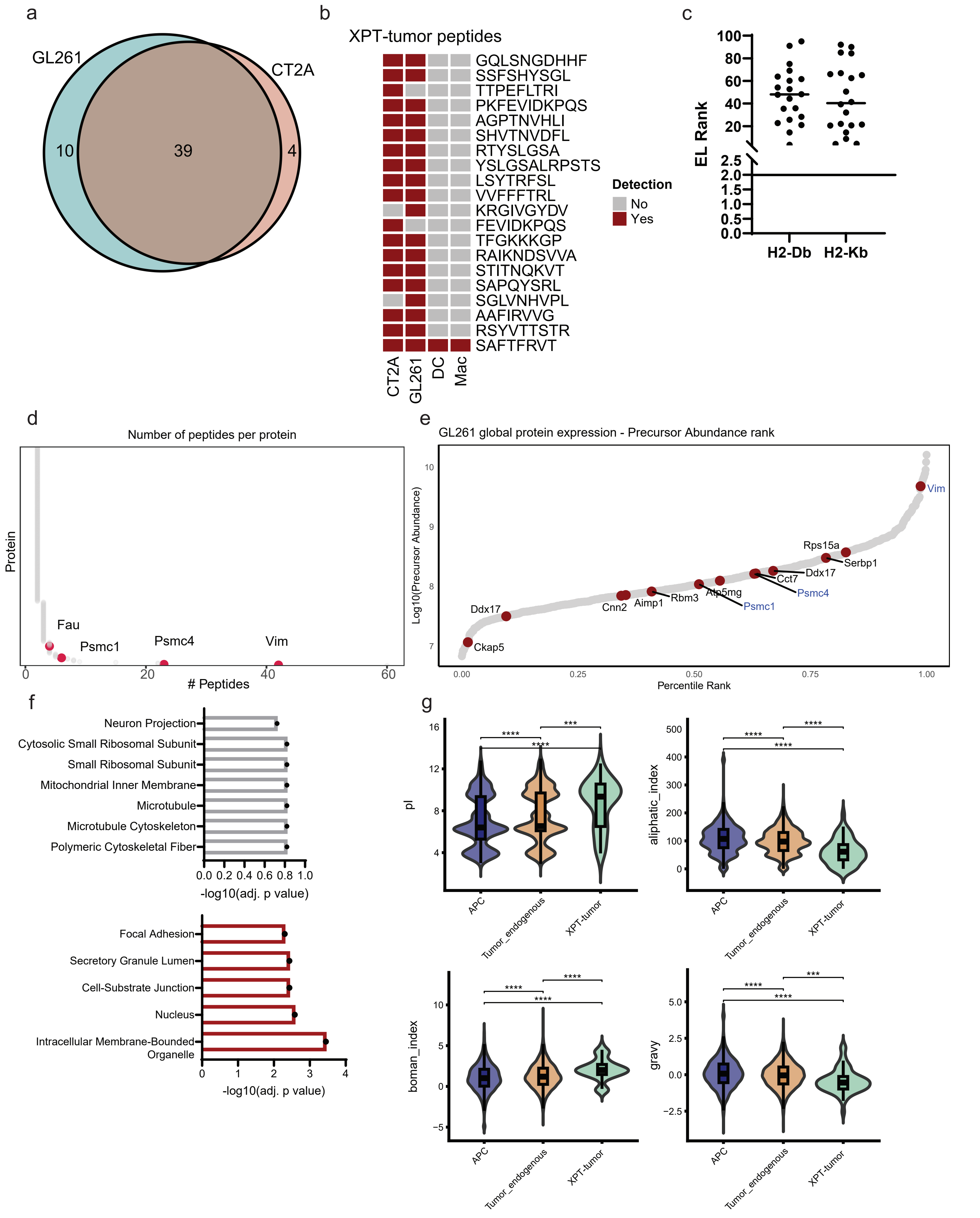

### Supplement 7

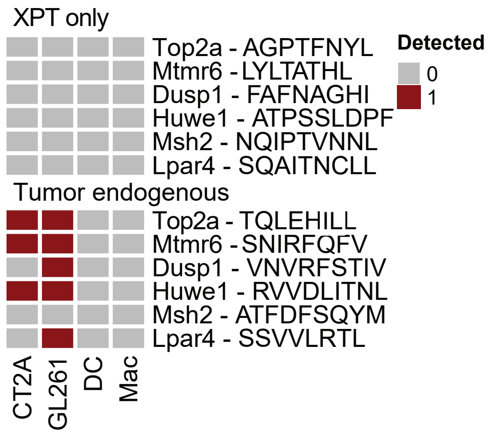

### Supplement 8

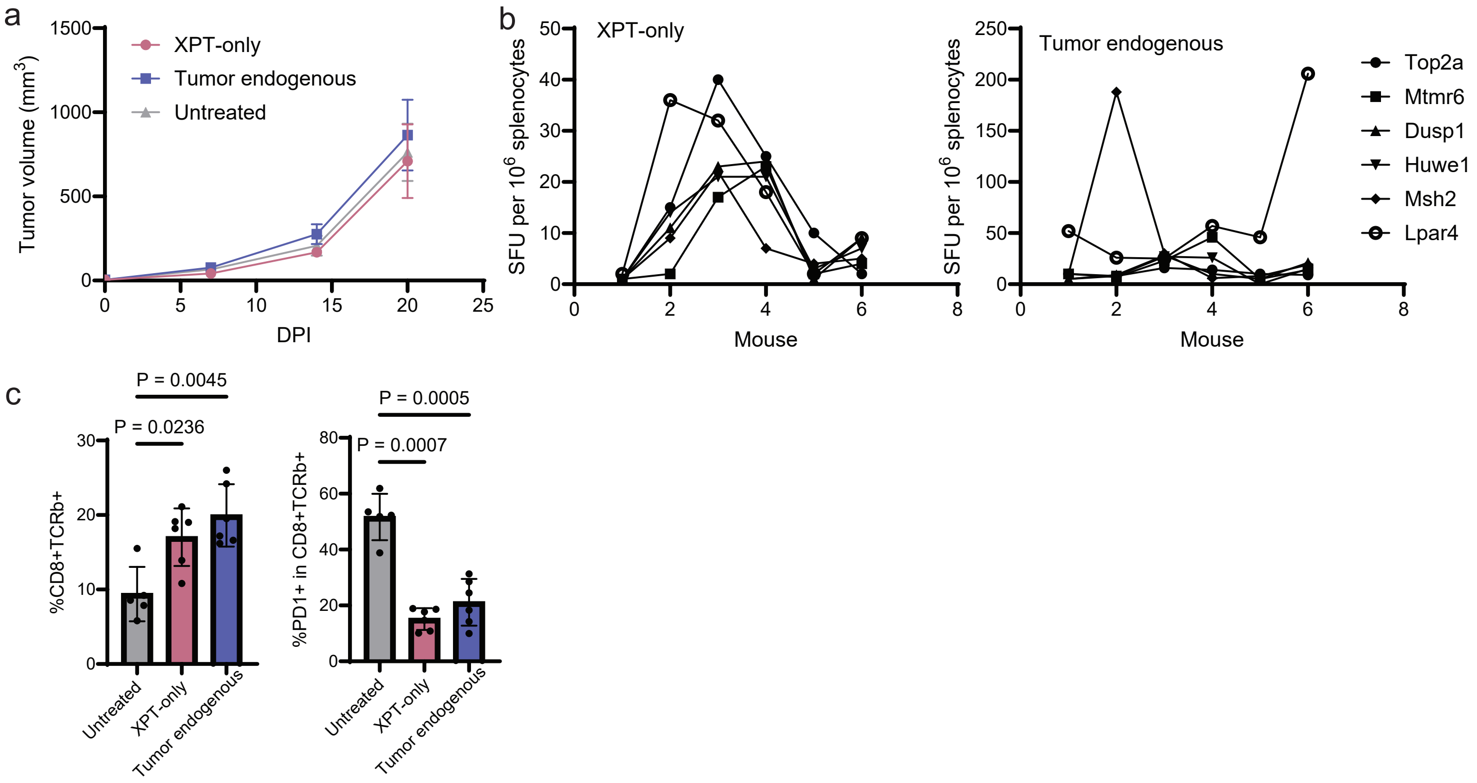

### Supplement 9

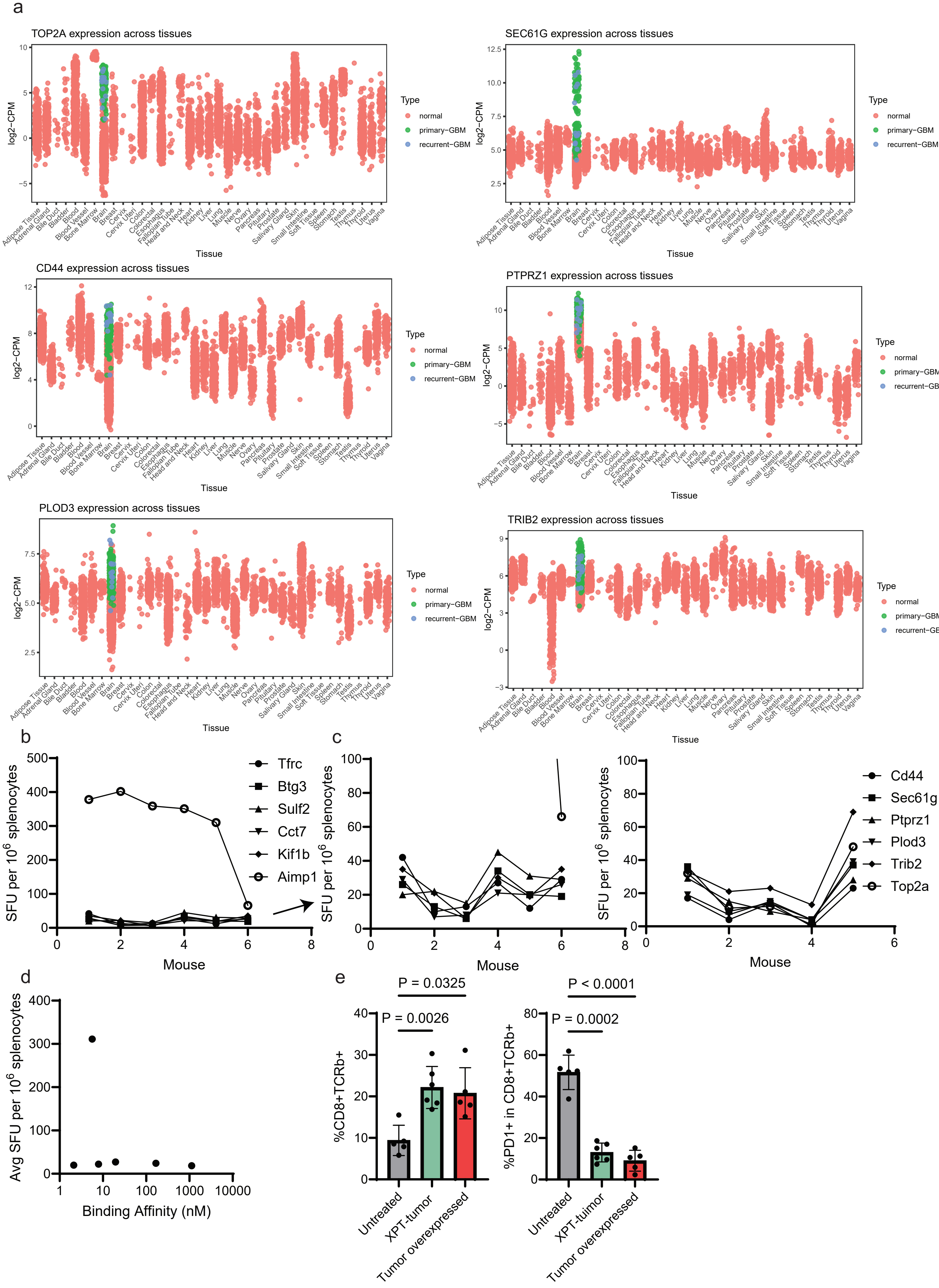
